# FG-Nup Sequence Length Polydispersity Enhances Selectivity of NPC Translocation

**DOI:** 10.1101/2025.02.07.637119

**Authors:** Manoj K Patel, Ajay S Panwar

## Abstract

The central channel of the nuclear pore complex (NPC) exhibits polydispersity in FG-nucleoporin (FG-Nup) sequence length, with longer FG-Nups on the periphery and shorter FG-Nups in the interior of the pore. The functional role of FG-Nup polydispersity in NPC transport was investigated using a minimal, coarse-grained model and Langevin dynamics simulations. The NPC was modelled as a cylindrical pore lined with a random copolymer brush composed of hydrophobic and hydrophilic segments, mimicking FG-Nups. The total number of hydrophobic segments corresponded to the fraction of FG-repeats, *f*, in the FG-Nups. The translocation of karyopherin-bound spherical cargo (12 nm spherical tracers) was simulated through two model NPCs; a homogeneous NPC (*h-*NPC) with uniform FG-Nup lengths and an inhomogeneous NPC (*ih-*NPC) featuring shorter FG-Nups in the middle and longer FG-Nups at the periphery. The *ih-*NPC demonstrated enhanced selectivity and significantly higher passage probabilities for karyopherin-bound tracers, with an increase of up to 90% compared to *h-*NPC. Analysis of binding contacts between tracers and FG-Nup hydrophobic segments revealed that tracer translocation was facilitated by a handover process between successive FG-Nups along the NPC length. The enhanced selectivity of the *ih-*NPC was attributed to an increase in binding contacts of the tracer with the shorter FG-Nups in its middle region. These findings provide a biophysical basis for the evolutionary significance of FG-Nup polydispersity in selective NPC transport.

## INTRODUCTION

The nuclear pore complex (NPC) is a large nanopore (internal diameter between 30 – 50 nm) on the nuclear envelope (NE) of a eukaryotic cell that regulates nucleocytoplasmic transport between the nucleus and the cytoplasm (1–5). It is one of the largest macromolecular assemblies in eukaryotic cells (1) with a highly conserved architecture and function across all eukaryotic species (3–5). Acting as a gatekeeper, the NPC regulates the bi-directional transport of proteins and nucleic acids across NE (1, 2), while permitting passive diffusion of small biomolecules and ions (< 40 kDa or < 6 nm) (6–8). Structurally, the NPC consists of protein subunits called nucleoporins (Nups), which are organized in an octagonal symmetry about the pore axis. Its central channel is lined with highly flexible, natively unfolded domains called FG-Nups, which constitute approximately 30% of the NPC mass (9–12). FG-Nups are characterized by repeats of hydrophobic phenylalanine (F) and glycine (G) residues that act as spacers between linker hydrophilic residues (13).

It is well established that transient hydrophobic interactions between hydrophobic domains in transport receptors or karyopherins (Kaps) and FG-repeats reduce the entry barriers for Kap-bound cargo. These interactions facilitate the efficient translocation of cargo complexes through the NPC and lead to the high degree of selectivity exhibited by the NPC (14–16). Experiments have confirmed that Kap-FG interactions are essential for facilitated transport through the NPC. However, the exact transport mechanism(s) are still under debate. In this context, several transport models of facilitated diffusion of Kap-bound cargo through the NPC have been proposed (12, 13, 17–21). Direct observation of FG-Nup dynamics and FG-Kap binding kinetics is challenging due to the complex organization of FG-Nups in the NPC and the heterogeneity in their sequences (22). Molecular simulation studies, primarily utilizing coarse-grained (CG) NPC models, have provided significant insight into several aspects of macromolecular transport through the NPC, including FG-Nup organization, FG-Nup/karyopherin binding and FG-Nup sequence heterogeneity. These simulations confirmed the key role of FG-Kap binding in reducing the energy barrier for entry into the NPC. However, previous studies have not examined the role of FG-Nup hydrophobicity on both FG-Nup organization and macromolecular transport through the NPC. Recently, we presented a polymer-based minimal CG model of FG-Nups to understand the effect of FG-Nup hydrophobicity on cargo translocation through the NPC (23). The model demonstrated the emergence of NPC selectivity and specificity over a narrow range of FG-Nup hydrophobic fraction, *f* (0.1 ≤ *f* ≤ 0.2). Distributions of hydrophobic FG-Nup fractions in yeast and human NPCs revealed that most hydrophobic fractions fall within the same range of *f*, suggesting a physical basis for this evolutionarily conserved feature (*f*) of NPCs.

Within the NPC, FG-Nups exhibit heterogeneity in FG-repeat groups and polydispersity in sequence length with respect to their location in the NPC. Among the core FG-repeat groups in yeast FG-Nups – such as FG, GLFG, F*x*FG, and LSFG (where *x* represents neutral hydrophilic amino acids) (24–26) – GLFG motifs are abundantly present in central FG-Nups. GLFG containing FG-Nups are known to form more cohesive interactions in comparison to the FG-Nups located in the peripheral NPC regions (27–29). FG-Nups always occur in multiples of eight, reflecting the underlying eight-fold symmetry of the NPC. Central FG-Nups are present in higher copy numbers compared to their peripheral cytoplasmic and nuclear counterparts. In addition, FG-Nups sequence lengths vary significantly along the pore axis (Figure 1A-1C). Whereas, central FG-Nups (such as Nup49, Nup57 and Nup145, 400 – 500 residues) exhibit shorter lengths, longer FG-Nups (such as Nsp1 and Nup145, 600 – 700 residues) tend to localize toward the peripheral regions of the NPC (24, 30–33).

**Figure 1.**
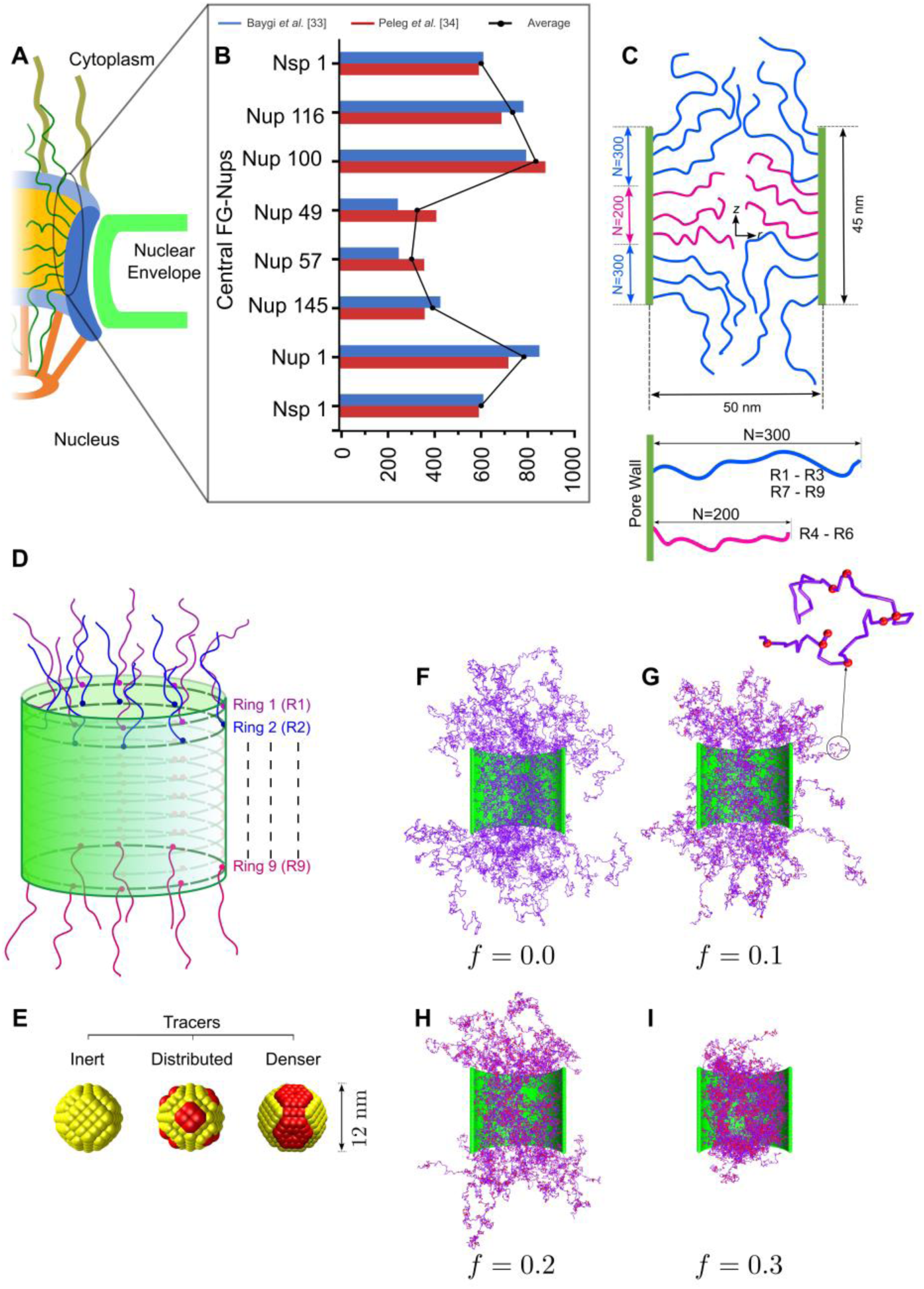
A minimal, coarse-grained model of the NPC and tracer particles. (A) Schematic of the NPC architecture showing, (B) measures and distribution of FG-Nup length along the NPC axis. (33, 34) (C) An inhomogeneous CG description of the NPC model incorporating FG-Nup length variations, with three rings of shorter FG-Nups (*N* = 200 beads) in the middle region and three rings each corresponding to longer FG-Nups (*N* = 300 beads) at the top and bottom peripheral regions of the NPC. (D) Rings are labelled from top to bottom by ring indexes, *R*1 to *R*9. (E, F, H, I) Side views of the equilibrated polymer brush for hydrophobic fractions *f* = 0.0 to *f* = 0.3 (G) Three types of tracers; inert (i-tracer), distributed patchy (sy-tracer), and denser patchy (hd-tracer).

These observations prompt questions on the functional significance of FG-Nup polydispersity (in sequence length) within the NPC. First, what is the underlying physical role of FG-Nup polydispersity with respect to NPC transport mechanisms and selectivity? Our minimal NPC model, which employed FG-Nups of uniform molecular weight, demonstrated that FG-Nup polydispersity is not essential for NPC selectivity (23). Does this suggest that NPCs may not strictly require FG-Nup sequence length polydispersity for selective transport? Second, would it not be more efficient for the NPC to adopt a simpler architecture, utilizing a single FG-Nup length or a more restricted sequence length distribution? Using a modified minimal NPC model (based on our previous NPC model that successfully demonstrated NPC selectivity) and Langevin dynamics simulations, the current study investigated the fundamental role of FG-Nup sequence length polydispersity in NPC transport. In contrast with our previous study, the modified model incorporated polydisperse FG-Nups with location-dependent sequence lengths, mimicking FG-Nup sequence length distribution along the NPC pore axis (see Figures 1A and 1B).

## SIMULATION METHOD

A minimal CG model was used to describe the NPC as a 45 nm long and 50 nm diameter cylindrical pore. Random copolymer brushes were terminally grafted to the inner periphery of the cylindrical pore to describe the natively unfolded domains of FG-Nups. Each random copolymer was a linear, bead-spring chain of *N* beads, with hydrophilic (*type 1*) and hydrophobic (*type 2,* each bead representing one FG repeat) beads randomly distributed along the chain. Hence, each polymer bead represents two adjacent residues in size. The number fraction of hydrophobic beads in an FG-Nup corresponded to the fraction of FG repeats, *f*, in the minimal NPC model. This CG model has been described previously in detail in our earlier work (23). However, the current setup differed from the previously reported model in the description of FG-Nup lengths along the pore axis. As shown in Figures 1C and 1D, nine rings of FG-Nups (identified by ring indexes, *R*1 to *R*9) were grafted on the inner periphery of the NPC. Each ring comprised 8 FG-Nups, thus reflecting the eightfold-symmetry of the NPC. In contrast to our previous work, FG-Nups of two different chain lengths were used to model the polydispersity in FG-Nup lengths observed in eukaryotic NPCs (Figures 1A and 1B). Whereas, the peripheral polymer brushes (top 3, *R*1 − *R*3 and bottom 3 rings, *R*7 − *R*9) were longer with *N* = 300 beads (modelled on the peripheral 600 residue Nsp1), the middle rings (*R*4 − *R*6) were shorter with *N* = 200 beads (modelled on the 400 residue Nup57) (Figures 1A and 1B) (33, 34). The bond length of the FG-Nup chain, *l*_0_, was chosen as the fundamental length scale in the simulation corresponding to a length of 1 nm which is approximately the size of two residues in a polypeptide chain. A detailed discussion of the choice of scales for length, force, time and energy can be found in our previous work (23). Thus, two types of NPCs were used in the current study; (i) *homogeneous NPC* (*h* −NPC) where *N* = 300 for all FG-Nups, and (ii) *inhomogeneous NPC* (*ih* −NPC) with shorter *N* = 200 FG-Nups grafted in the middle rings (see Figures 1C and 1D).

Actively diffusing cargo particles, representing proteins moving across NE, were modeled as hollow, rigid spheres with a tracer diameter, *d*_*t*_ = 12 nm. Three types of tracers were considered; (i) inert tracers (*i-*tracer) without any FG-binding domains, and tracers with FG-binding domains including (ii) distributed patchy tracers (*sy-*tracer), and (iii) denser patchy tracer (*hd-*tracer). A shown in Figure 1G, each tracer was composed of the 276 CG beads of two types; type 5 which were inert (depicted as yellow), and type 6 (representing Kap binding domains, red patch on yellow surface). The type 6 beads interacted with the type 2 beads (red) on FG-Nups through an attractive Lennard-Jones interaction, that modelled affinity between FG-repeats and Kaps. Although both *sy-*tracer and *hd-*tracer contained the same number of type 6 beads, they differed in how these beads were distributed on their respective surfaces. Whereas, FG-binding domains (patches of type 6 beads) were uniformly distributed for the *sy-*tracer, they were clustered into a large, localized patch on the *hd-*tracer. Notably, the *hd-*tracer description models the karyopherin-bound cargo more closely because FG-binding domains on Kaps (including importin-β, Kap95) are asymmetrically positioned on their surface (6, 35–38). This is essential for cargo recognition by FG-Nups and subsequent entry and translocation through the NPC.

Implicit solvent Langevin dynamics simulations were carried out to simulate tracer translocation through both *h-*NPC and *ih-*NPC for FG hydrophobic fractions in the range, 0 ≤ *f* ≤ 0.4. The equations of motion were integrated in the NVE ensemble using the molecular dynamics package, LAMMPS (39, 40). A Langevin piston maintained the temperature at 310 K. The system (NPC and tracer) was placed in a cuboidal simulation box of volume 100 × 100 × 250*l*_*o*_^3^, with fixed boundaries. The cylindrical pore spans from −22.5*l*_*o*_ ≤ *z* ≤ 22.5*l*_*o*_ in the axial z-direction. The dimensionless Lennard-Jones (LJ) interaction energies between hydrophobic (ɛ_22_ = 1.5) and hydrophilic ɛ_11_ = 0.1 beads were chosen to reflect accurate scaling behavior for homopolymers under different conditions (41). In addition, and interaction energy ɛ_26_ = 2.0 was used to model the affinity between FG-repeats and Kaps. Tracers were initially placed at *z* = 90 *l*_0_, far from the extended polymer brushes of the NPC (see Figure 1E). In the NPC, directionality of macromolecular translocation is determined by a Ran gradient (42) across NE. The is modelled by a downward force, *F*_*t*_ = 2 pN, acting on the tracer (25, 33, 43–46) throughout the translocation simulation. *F*_*t*_ is an over-simplified representation of a more complex process, and could also be attributed to electrostatic interaction between negatively charged importin and positively charged FG-Nups (majorly along pore axis) which could favor transport rate (47–49). The adoption of a constant downward force, *F*_*t*_ captures key aspects of cargo motion in the biological system. The reader is referred to our previous work for a detailed justification of energy scales used in the current work (23).

## RESULTS AND DISCUSSION

### Tracer Trajectories

Tracer translocation was simulated through both homogeneous (*h-*NPC) and inhomogeneous (*ih-* NPC) pores. As detailed in the Methods section, three types of tracers were simulated for translocation through the NPCs: inert (*i-*tracer), distributed (patchy, *sy-*tracer) and dense (patchy, *hd-*tracer). A constant downward force of *F*_*t*_ = 2 pN was applied on the tracer during translocation. Figure 2 summarizes the comparison of tracer trajectories for the three tracer types across the two types of NPCs at an FG fraction, *f* = 0.2. Trajectories for all tracer types corresponding to hydrophobic fractions, 0 ≤ *f* ≤ 0.4, are shown in Figures S1–S3 (*h-*NPC) and Figures S4–S6 (*ih-* NPC) of the Supplementary Information. Previously, we have shown that NPC selectivity emerged for an FG-fraction range of 0.05 ≤ *f* ≤ 0.2 in an *h-*NPC, corresponding to hydrophobic fractions observed in both yeast and human FG-Nups (23). Therefore, the results for *f* = 0.2, highlighted in Figure 2, are representative of a minimal pore that mimics NPC selectivity. However, tracer trajectories corresponding to the entire hydrophobic fraction range (0 ≤ *f* ≤ 0.4) were simulated for the sake of completeness. A total of twenty trajectories were simulated for each case. Based on their simulation outcome, trajectories were categorized as rejected (black), trapped (magenta) or translocated (blue), as shown in Figure 2 (and Figures S1-S6 of the Supplementary Information).

**Figure 2.**
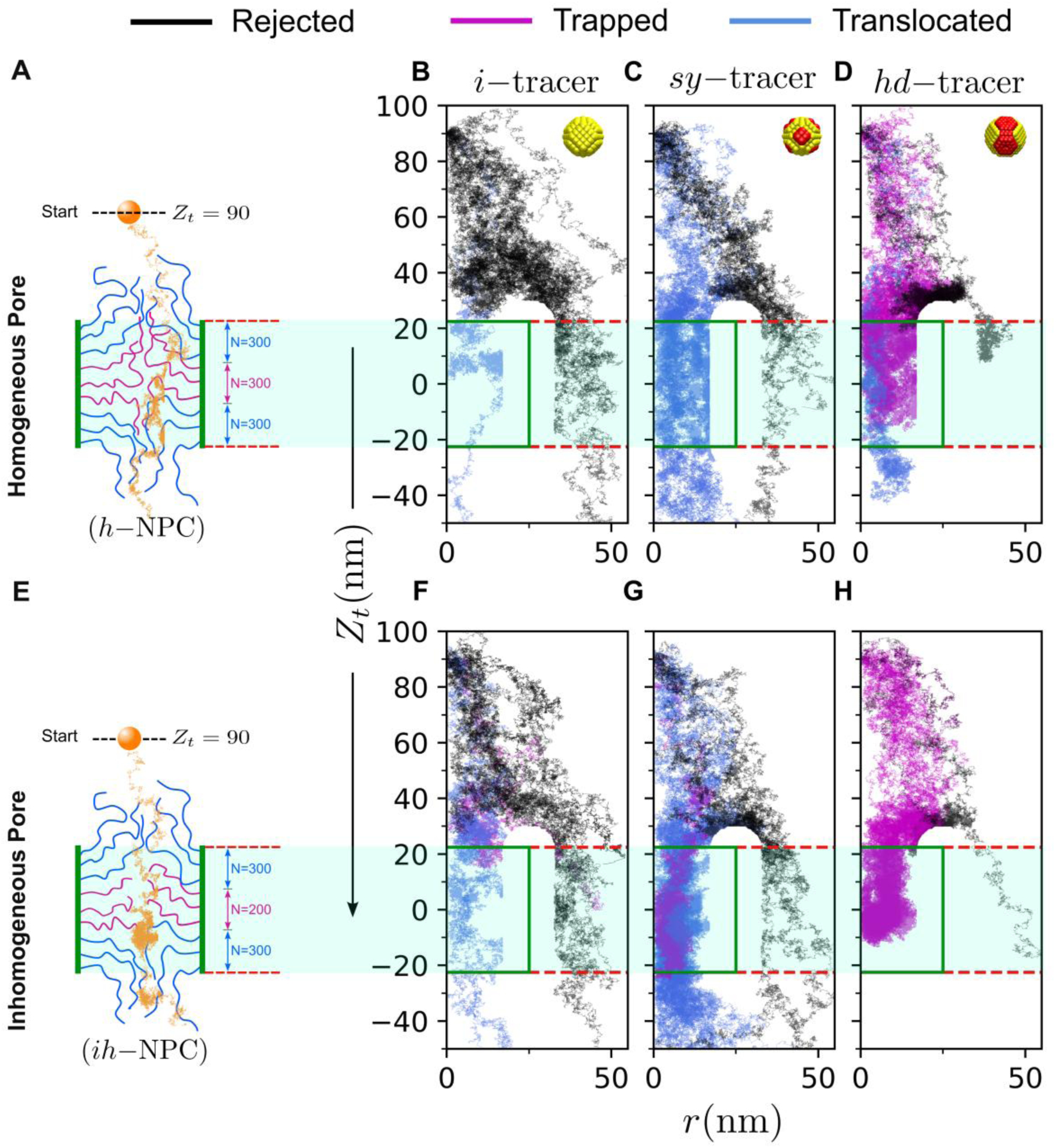
Tracer trajectories from twenty independent simulations shown for *f* = 0.2 corresponding to (A, F) *i-*tracer, (B, G) *sy-*tracer and (C, H) *hd-*tracer, respectively. The trajectories show the variation of the *Z*-coordinate of the tracer, *Z*_*t*_, with respect to the tracer radial coordinate, *r*. The cylindrical NPC is shown by a green rectangle extending from − 22.5 ≤ *z* ≤ 22.5. The simulated trajectories were classified into three types, successful (blue), trapped (pink) and rejected (black).

The trajectories shown in Figures 2B and 2F clearly show that both *h-*NPC and *ih-*NPC almost entirely rejected inert tracers (cargoes without Kaps). However, a very small number of inert trajectories (1 – 3 out of 20) translocated through the pores, a result attributed to artifacts of our CG model, as previously discussed in Patel *et al.* (23). An increase in the density of patches from the distributed patchy to the denser patchy tracer, resulted in more trajectories entering the NPCs (or fewer rejected trajectories). This was true for both *h-*NPC (Figure 2C *versus* Figure 2D) and *ih-*NPC (Figure 2G *versus* Figure 2H). FG-Nups are known to form an extended brush outside the pore that presents an entropic barrier to tracer entry into the NPC (8, 12, 17, 23). Simultaneously, these extended FG-Nups serve as docking sites for Kap-bound cargo (patchy tracers) through hydrophobic interactions between FG repeats and Kap binding domains (6, 50). These hydrophobic interactions reduce the entropic penalty, thereby facilitating cargo entry into the NPC. A visual comparison of patchy tracer trajectories indicated that the *ih-*NPC allowed more trajectories when compared to the *h-*NPC. This seemed to be true for both *sy-*tracer and *hd-*tracer. However, a higher number of trajectories were trapped inside the *ih* −NPC compared to the *h-* NPC. For example, none of the *hd-*tracer trajectories that entered the *ih-*NPC successfully translocated through the pore. The following analysis below quantifies tracer translocation through both types of NPCs to understand the effect of FG-Nup polydispersity in *ih-*NPC.

### Pore response to different types of tracers

The interaction of the tracer with FG-Nups and its translocation through the NPC was analyzed as a three-stage process, including, (i) entry into the pore, (ii) arrival at the mid-plane and (iii) exit from the pore. Three separate probabilities for entry, *p*_*e*_, crossing the mid-plane, *p*_*m*_, and successful passage of the tracer, *p*_*p*_ were calculated for every case. The planes *z* = 22, *z* = 0 and *z* = −22 were used as thresholds to define *p*_*e*_, *p*_*m*_ and *p*_*p*_, respectively, in the following way,

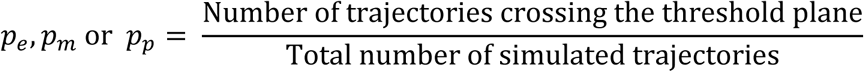

Figure 3 shows that the entry of inert tracers was almost entirely blocked for both *h-*NPC and *ih-* NPC (*p*_*e*_ ≈ 0.2). The unusually high values of *p*_*e*_ observed for *f* = 0.4 should not be interpreted as tracer entry into the pore. In fact, tracer transit was blocked by a shallow crater-like surface formed due the complete collapse of the FG-Nup brush for *f* ≥ 0.25 (see Figure S7-S10 and segment density discussion in our previous work (23)). The collapsed brush resulted in a high segment density inside the pore which effectively sealed the pore. In the physically relevant range (0.05 ≤ *f* ≤ 0.2), *p*_*e*_, *p*_*m*_ and *p*_*p*_ values were nearly the same for inert tracers passing through both *h-*NPC and *ih-*NPC, indicating that all trajectories that entered the pore successfully passed through the pore. However, as indicated above, the successful translocation of an inert tracer trajectory is an artefact of our CG model (23). Hence, Figures 3A-C indicate that inert tracers were effectively rejected by both *h-*NPC and *ih-*NPC.

**Figure 3.**
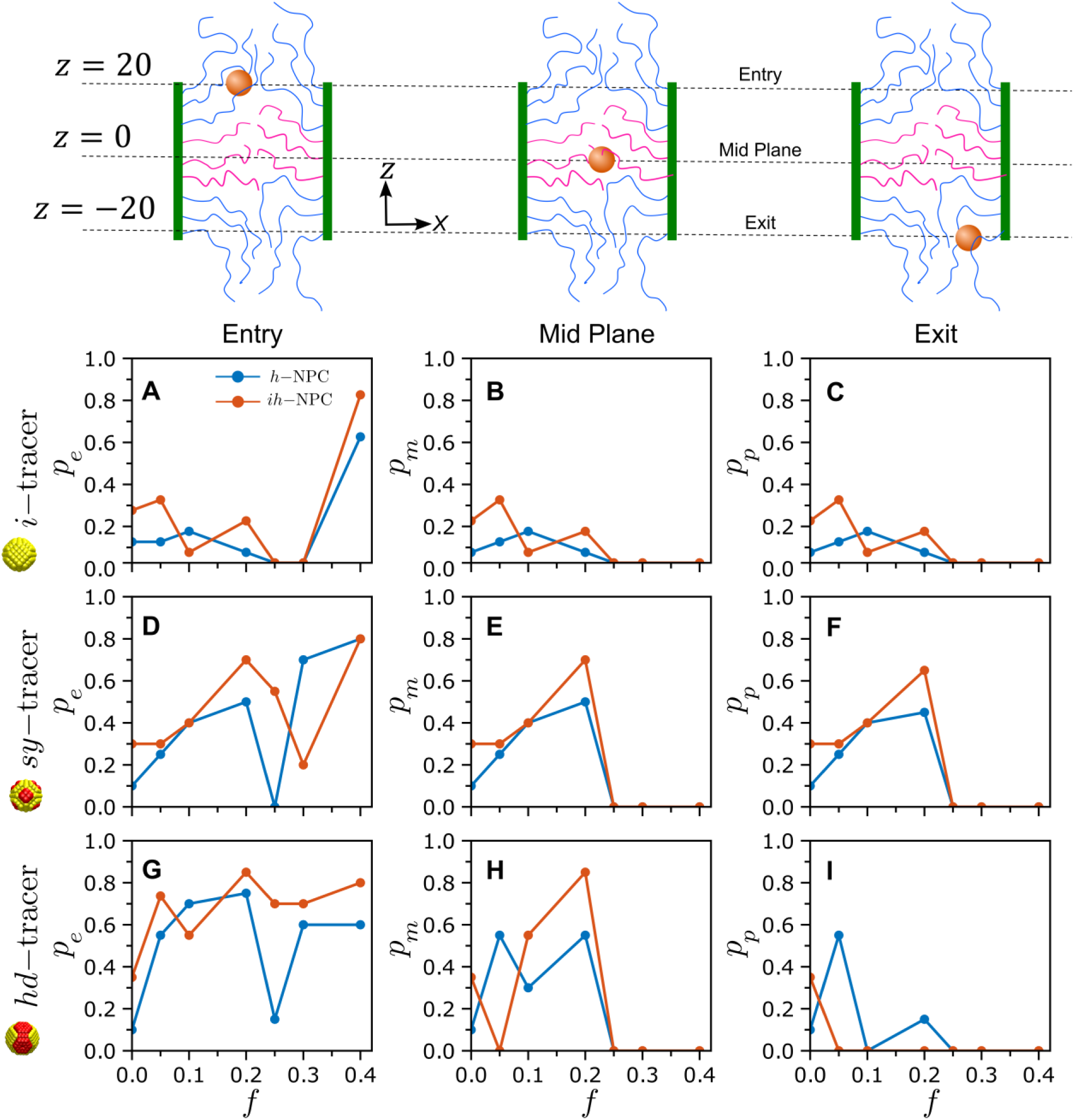
Plots showing different probabilities associated with tracer translocation as function of *f*, including the probability of tracer entry, *pe*, probability of crossing the NPC mid-plane, *pm*, and the probability of the tracer completely exiting the pore, *pp*. Plots are shown for i-tracer (A – C), sy-tracer (D – F) and hd-tracer (G – I) corresponding to both h-NPC and ih-NPC.

In contrast to inert tracers, distributed patchy tracers showed significantly higher values for *p*_*e*_, *p*_*m*_ and *p*_*p*_. No significant changes in *p*_*e*_, *p*_*m*_ and *p*_*p*_ values were observed as the *sy-*tracer translocated through the pore (Figures 3D–F). In other words, the probability plots indicated no trapping of tracers during transit through the pore. However, probability distributions were narrower and higher for distributed patchy tracer translocation through *ih-*NPC in comparison to *h-*NPC. For *f* = 0.2, a 56% increase in *p*_*p*_(from 0.45 to 0.65) was observed for peak values for the *ih-*NPC. Previously, our minimal *h-*NPC model showed that both NPC selectivity and specificity emerged in the biophysically relevant range 0.05 ≤ *f* ≤ 0.2 (23). A comparison of *p*_*p*_ distributions in Figure 3F indicated that the polydisperse FG-Nup description of *ih-*NPC resulted in a more selective NPC with a significantly higher probability of successful translocation for the same range of *f*.

The localization of hydrophobic patches in *hd-*tracer resulted in large *p*_*e*_ values, between 0.6 – 0.9, in the relevant range (disregarding *f* ≥ 0.25 in Figure 3G). This was due to increased binding of FG repeats to the large patch on the tracer. Increased tracer-FG repeat binding also increased tracer access to the pore midplane (*p*_*m*_ values in Figure 3H). Figure 3H shows that peak *p*_*m*_ values (at *f* = 0.2) for *ih-*NPC were almost 50% higher when compared to *h-*NPC – reinforcing the idea that polydispersity in FG-Nup sizes improved tracer access to the NPC. A comparison of Figures 3E and 3H showed that *hd-*tracer demonstrated higher *p*_*m*_ values than the *sy-*tracer for *ih-*NPCs. However, the localization of all hydrophobic patches in the *hd-*tracer lead to significant trapping of tracers within the NPCs. Tracers were completely trapped for all *f* (except *f* = 0) for the *ih-* NPC (Figure 3I). Despite a significant difference between *p*_*m*_ and *p*_*e*_ values, the *h-*NPC did allow limited passage for denser patchy tracer trajectories (Figure 3I).

### Ran Gradient: Impart directionality and dissociate Kap-cargo-complex

The Ran-GTP concentration gradient between the cytoplasm and nucleus ensures directionality for Kap-bound cargoes during nucleocytoplasmic transport (6, 43). The nucleus contains GTP-bound Ran, while GDP-bound Ran resides in the cytoplasm (43, 44, 51).Upon nuclear entry of Kap-bound cargo, RanGTP promotes cargo dissociation from importins by triggering conformational changes that reduce the Kap’s affinity for the cargo (52–54). These have an additional effect of lowering FG-repeat binding affinity to the Kaps (20, 53, 55–60). Conversely, during export, exportins transport RanGTP-associated cargo to the cytoplasm, where GTP hydrolysis releases the complex (55, 61).

In our simulations, the RanGTP-dependent cargo-complex dissociation was mimicked by weakening the hydrophobic interactions between the *hd-*tracer and the middle FG-Nup rings belonging to both *h-*NPC and *ih-*NPC. This was achieved by decreasing the LJ interaction energy between FG-repeats and tracer hydrophobic patches from an initial value of ɛ_26_ = 2.0 to ɛ_26_ = 1.5. It should be noted that the change was applied only to the hydrophobic beads belonging to the middle FG-Nup rings (*R*4 – *R*6, see Figure 1). The interaction energy values for outer rings (*R*1 – *R*3 and *R*7 – *R*9) remained unchanged at ɛ_26_ = 2.0. The change in affinity is illustrated through the “before” and “after” schematics in Figures 4A and 4B, respectively. The impact of the reduced affinity is evident in the “before” and “after” plots of tracer trajectories (*f* = 0.1, 0.2 for *ih-*NPC) in Figures 4A and 4B. Notably, nearly all trapped trajectories (majenta) were successfully released (blue) by the *ih-*NPC (and for translocations through *h-*NPC, see Figure S11-S12).

**Figure 4.**
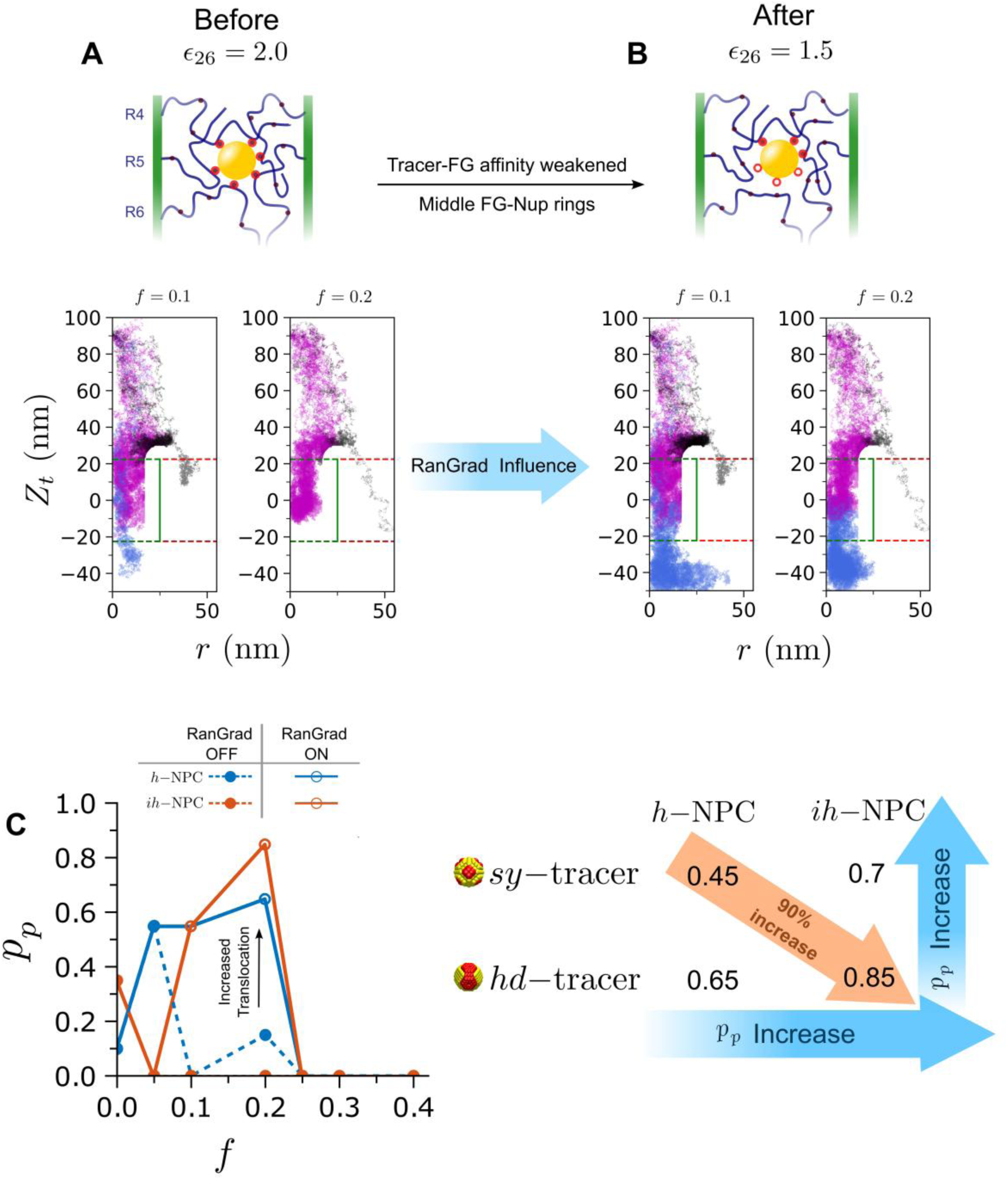
Lowering the interaction strength, *∈*_26_, between the tracer and middle FG-Nup rings enabled the release of trapped *hd*-tracers in both *h*-NPC and *ih*-NPC. Trajectories shifted from being trapped (A, magenta) to successfully exiting the NPC (B, blue) following the reduction in interaction strength. (C) Comparing *p_p_* values between *h*-NPC and *ih*-NPC, before and after reducing *∈*_26_. The value of *p_p_* for hd-tracer in *ih*-NPC was nearly 90% more than the value for *sy*-tracer in *h*-NPC.

The high *p*_*m*_ and low *p*_*p*_ values for *hd-*tracer in Figures 3H and 3I, respectively, imply that most tracers were trapped at the mid-plane. The trapping is likely due to the strong binding between FG-repeats from *R*4 – *R*6 (grafted at the mid-plane) and hydrophobic patches of the *hd-*tracer, as observed in the trajectories in Figure 3I. This observation formed the rationale for selectively decreasing ɛ_26_ value to 1.5 for *R*4 – *R*6. It should be noted that ɛ_26_ = 1.5 defines the FG-FG interaction level in our model. Consequently, the reduced patch-FG binding energy is as strong as the hydrophobic interactions between FG-Nups inside the NPC.

Figure 4C shows the change in *p*_*p*_ values following the reduction in ɛ_26_ value to 1.5 for *R*4 – *R*6. In the biophysically relevant range, 0.1 ≤ *f* ≤ 0.2, the release of trapped trajectories resulted in a dramatic increase in the probability of successful translocation (*p*_*p*_ values between 0.6 to 0.9) for both *h-*NPC and *ih-*NPC. At their maximum, *p*_*p*_ values were about 30% higher for *ih-*NPC when compared with *h-*NPC. For instance, *p*_*p*_ ≈ 90% at *f* = 0.2 for *ih-*NPC indicating successful translocation for almost all simulated trajectories. A comparison of the *p*_*p*_ distributions for *h-*NPC and *ih-*NPC (solid blue and red curves in Figure 4C, respectively) shows that the distribution is narrower and exhibits higher values in the case of *ih-*NPC, following the reduction in ɛ_26_. Together, these two attributes of *p*_*p*_ distribution indicate that the *ih-*NPC is a more selective pore in comparison to the *h-*NPC (see Figure S13).

Although, the *h-*NPC model can generate NPC selectivity for 0.05 ≤ *f* ≤ 0.2 (23), polydispersity in FG-Nup sizes (*ih-*NPC) enhances the selectivity of the NPC (Figure 4A). Notably, the FG-Nup size distribution used in the *ih-*NPC case mimicked the FG-Nup sequence lengths reported for yeast NPCs, as shown in the model setup in Figures 1B and 1C. Hence, the result provides a biophysical reason for the variation of FG-Nup sequence lengths along the NPC axis (33, 34). In addition, *p*_*p*_ values for *ih-*NPC were higher for the *hd-*tracer in comparison to the distributed patchy tracer. This result is consistent with the *hd-*tracer being a more realistic mimic of karyopherin-bound cargo, which is characterized by localized placement of FG-binding domains (6, 35, 38, 53, 62). The physical mechanisms underlying selectivity enhancement in the *ih-*NPC are not immediately evident from the results of Figures 3 and 4. Polydispersity in FG-Nup sizes can influence interactions between FG-Nups and tracers, as well as interactions among FG-Nups themselves. These interactions can alter both tracer binding to FG-Nups and the brush morphology inside the *ih-*NPC.

### Mesh: polymer network crosslinks and trapped trajectories

In our previous work (23), we proposed a hybrid NPC transport model that combined effects of entropic repulsion from FG-Nups and size exclusion due to the formation of a sieve-like network inside the NPC. Hence, our model exhibited features of both entropic repulsion model (17) and the selective-phase model of NPC transport (63). This hybrid approach also aligns with qualitative *in vitro* characterizations of individual FG-Nups (13, 55, 64), though consensus on the *in vivo* scenario remains elusive (65). In our earlier analysis of the *h-*NPC (23), a single characteristic mesh size, *ξ*, was calculated for the entire pore. However, in the *ih-*NPC, variations in FG-Nup size at different regions within the pore may lead to region-dependent mesh sizes. A variation in *ξ* may be responsible for enhanced values of *p*_*p*_ in the *ih-*NPC.

To quantify the mesh size, the central channel was divided into three regions: *top* (7.5 ≤ *l*_*t*_ ≤ 20), *middle* (−7.5 ≤ *l*_*t*_ ≤ 7.5) and *bottom* (−20 ≤ *l*_*t*_ ≤ −7.5), as shown in Figures 5A – 5D. The average mesh size ⟨*ξ*⟩ of the equilibrated brush (averaged over the last 1000 simulation frames) was calculated using the Stillinger algorithm (66, 67) as implemented in our previous work (23). Surprisingly, the mesh size values, *ξ*, were nearly the same across all three regions. In addition, *ξ* values were also nearly the same for both *h-*NPC and *ih-*NPC. It is interesting to note that mesh formation was not observed for *f* < 0.15 near the entry and exit regions of the pore (*top* and *bottom* in Figures 5B and 5D, respectively). This was true for both types of NPCs and can be attributed to greater chain entropy at the pore openings. In all three regions, *ξ* decreased continuously with increasing *f* and dropped below 6 nm for *f* > 0.2. This resulted in size exclusion of the *d*_*t*_ = 12 nm tracers (rejection by the NPCs) for all FG hydrophobic fraction, *f* > 0.2, which was consistent with our previous findings (23). However, absence of heterogeneities in *ξ* indicates that mesh size does not determine the enhanced selectivity observed in the *ih-*NPC.

**Figure 5.**
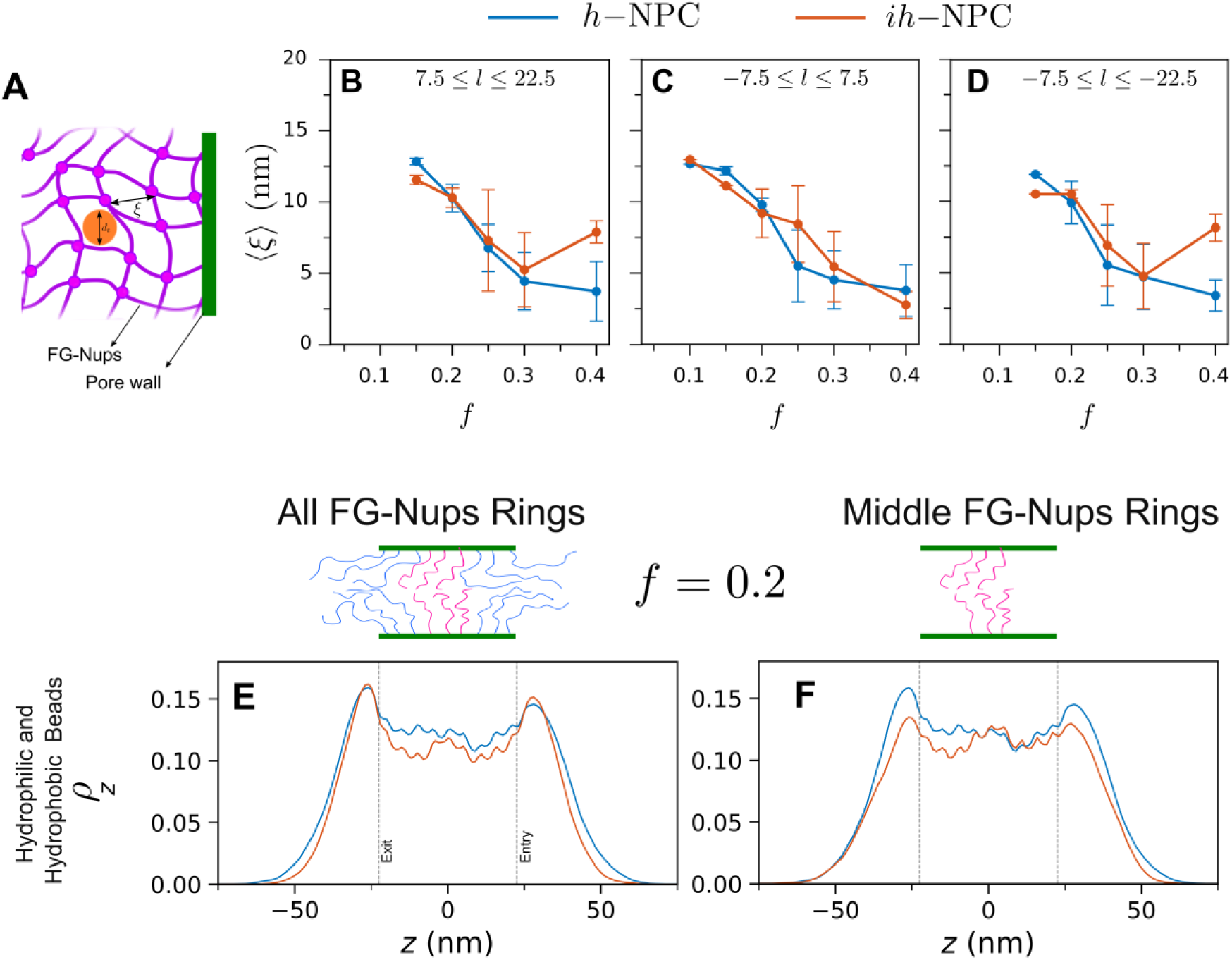
Average mesh size measured across three distinct zones within the pore: top (A), middle (B), and bottom (C). No mesh formation was observed in the outer region of the pore for (*f* <0.15). Plots of *ρ*_*z*_ of FG-Nups for *f* = 0.2 corresponding to, (E and F) all hydrophilic and hydrophobic bead. A distinction is also made for *ρ*_*z*_ values for all rings, *R*1 − *R*9 (E) and only for the middle rings, *R*4 − *R*6 (F).

Figures 5E and 5F plot the FG-Nup axial densities, *ρ*_*z*_, for *f* = 0.2 in both *h-*NPC and *ih-*NPC. These values are calculated by taking both hydrophilic and hydrophobic beads of FG-Nups into account and averaged over the final 100 frames of the equilibrated brush trajectories. Whereas, *ρ*_*z*_ values in Figures 5E account for FG-Nup segments from all nine rings, *R*1 – *R*9, *ρ*_*z*_ values plotted in Figures 5F were calculated only for the middle rings *R*4 – *R*6. Similarly, *ρ*_*z*_ plots for all values of *f* are shown in the Supplementary section, Figures S7 – S10. Axial density profiles appeared to be very similar for the two types of NPCs when *ρ*_*z*_ is averaged over all FG-Nups (Figure 5E). However, the differences between the *h-*NPC and *ih-*NPC are highlighted when *ρ*_*z*_ is plotted for the middle ring FG-Nups. Plots in Figures 5F show significantly larger *ρ*_*z*_ values at the pore openings in the case of *h-*NPC. For the *h-*NPC, the larger FG-Nups from the middle rings (300 beads/ 600 residues) were more extensively expelled from the pore interior due to entropic confinement. Conversely, the shorter middle ring FG-Nups in the *ih-*NPC (200 beads/ 400 residues) experienced a lower entropic penalty, leading to smaller *ρ*_*z*_ values at the pore openings. This was confirmed by an increase in *ρ*_*z*_ near the mid-plane for *ih-*NPC (near −7 < *z* < 7). Entropic repulsion due to extended FG-Nup brush conformations outside the NPC forms the basis of the “virtual gate model” (17). This effect was effectively demonstrated in the near complete rejection of inert cargoes in our previous study using an *h-*NPC (23). A more extended brush conformation increases tracer repulsion, thereby enhancing the likelihood of the tracer being rejected by the pore. The middle ring FG-Nups in the *h-*NPC exhibited higher *ρ*_*z*_ values at the pore openings, potentially providing greater entropic repulsion to the incoming tracer. Additionally, differences in *ρ*_*z*_ values between *h-*NPC and *ih-*NPC may influence tracer binding to FG-Nups and its subsequent transport through pore.

### FG-Nup rings: recognition and handover of tracer

Irrespective of the pore type, tracers moving into the pore first encounter the peripheral FG-Nup rings (*R*1– *R*3) at the pore entry, before interacting with the middle FG-Nup rings (*R*4– *R*6, see the ring arrangement in Figure 1). Differences in *ρ*_*z*_ for middle ring FG-Nups between *h-*NPC and *ih-*NPC (discussed in Figure 5G and 5H) may influence how tracers interact and bind with successive rings during the translocation process. The interactions between the tracer and FG-Nups belonging to a specific ring were estimated as the tracer moved through the pore. For a given axial location of the tracer, *Z*_*t*_, this was achieved by calculating the average number of binding contacts, *n*_*b*_, between the hydrophobic patches on a tracer and FG-repeats for a specific ring. A distance cut-off of 1.5 nm between the FG-repeats (hydrophobic FG-Nup beads) and patchy beads of a tracer was used to determine a binding contact. For each position, the average was taken over 200 frames which comprises 100 frames before and after the specific *Z*_*t*_ location. Figure 6 illustrates tracer binding to various rings at a specific axial location, *Z*_*t*_, by plotting *n*_*b*_(*Y*-axis) as a function of the ring index, *R* (*X-*axis). The ring index, *R*, takes integer values ranging from 1 to 9 corresponding to rings, *R*1 through *R*9, respectively. Plots are shown for both the *hd-* tracer (Figure 6A) and for the *sy-*tracer (Figure 6B), respectively, corresponding to *f* = 0.2. For each tracer type, *n*_*b*_ *versus* ring index is shown at three tracer locations, (i) *entry* (*Z*_*t*_ = 20), (ii) *mid-plane* (*Z*_*t*_ = 0), and (iii) *exit* (*Z*_*t*_ = −20). Further, these plots compare the binding interactions between *h-*NPC and *ih-*NPC for each case. Similar plots for *f* = 0.1 are included in the Supplementary Information Figure S14.

**Figure 6.**
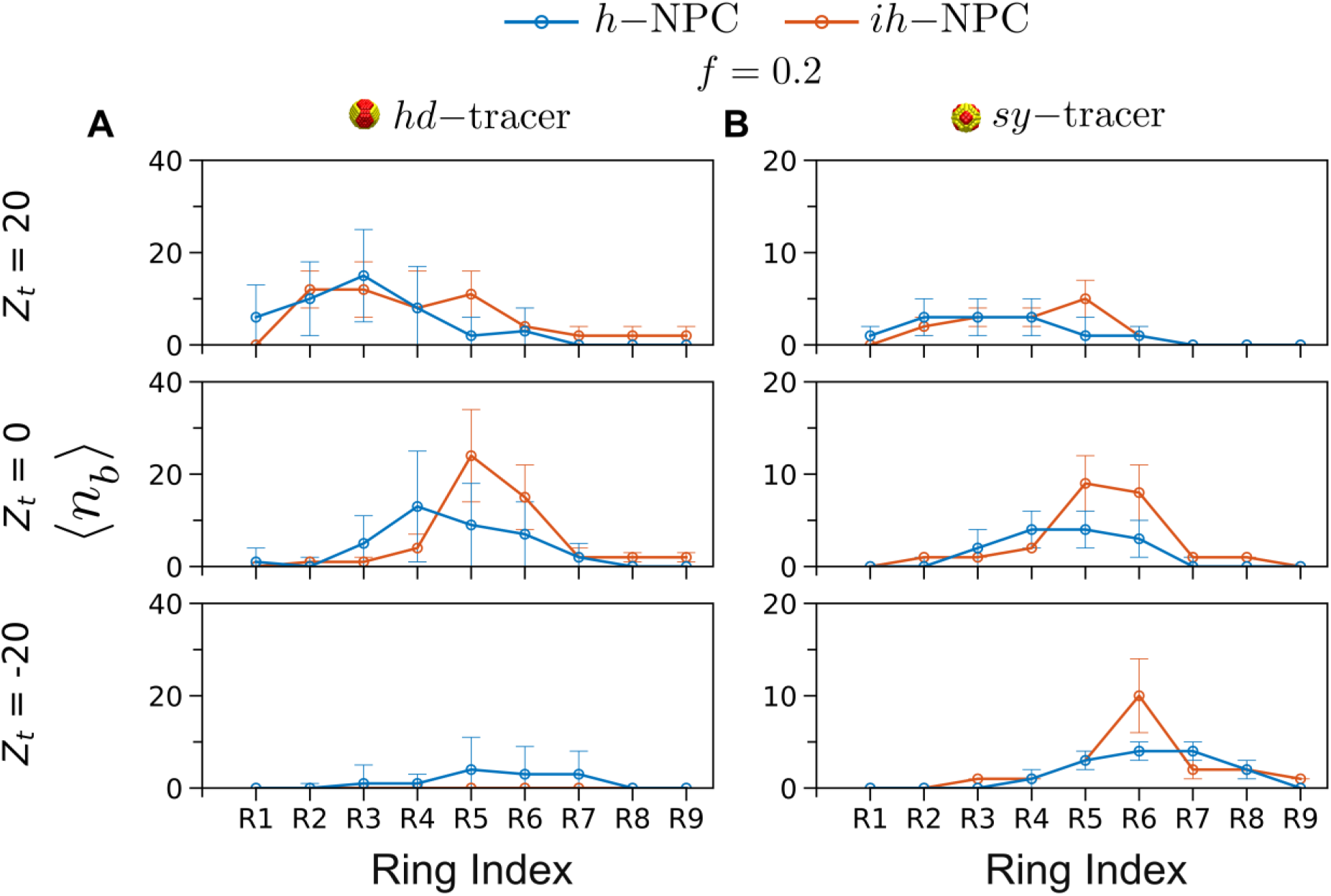
Average number of binding contacts, 〈*nb*〉, between tracer binding domains and FG-repeats corresponding to individual rings, *R*1 − *R*9, shown for both *h* −NPC and *ih* −NPC. The variation in 〈*nb*〉 with respect to the ring index are shown at the entry (*Zt* = 20), mid-plane (*Zt* = 0) and exit (*Zt* = −20) of the NPCs for (A) hd-tracer and (B) sy-tracer.

Figures 6A and 6B (and Figure S14) show that *n*_*b*_ values for the *hd-*tracer exceeded corresponding values for the *sy-*tracer by more than two times. This clearly demonstrated that a higher density of FG-binding contacts resulted in increased FG-Nup binding to the tracer. For the *hd-*tracer at the pore entrance (*Z*_*t*_ = 20), the distributions of binding contacts with different rings differed between *h-*NPC and *ih-*NPC. In the case of the *h-*NPC, the tracer was bound largely to peripheral rings *R*1 – *R*3. However, in the case of the *ih* −NPC, middle rings *R*4 – *R*6 also formed significant binding contacts with the *hd-*tracer. As a result, a shift in the *n*_*b*_ plot toward *R*4 – *R*6 was observed at *Z*_*t*_ = 20 for the *ih-*NPC. This was direct consequence of the difference in *ρ*_*z*_ values for middle ring FG-Nups between *h-*NPC and *ih*-NPC (Figure 5F). Whereas, the longer FG-Nups in *h-*NPC stretched outside the pore, the shorter FG-Nups in *ih-*NPC were present in greater fraction inside the pore resulting in higher *n*_*b*_ values in *ih-*NPC.

The difference in *n*_*b*_ plots between the *h-*NPC and *ih-*NPC became more pronounced when the tracer moved to the mid-plane (*Z*_*t*_ = 0). The tracer formed significantly more contacts with the middle rings in *ih-*NPC in comparison to *h-*NPC. Whereas, the peak in *n*_*b*_ was observed at *R*4 for *h-*NPC, it appeared at *R*5 for the *ih-*NPC. For both *h-*NPC and *ih-*NPC, the *n*_*b*_ plots showed a shift in their respective peak values from outer rings (*R*1 – *R*3) at *Z*_*t*_ = 20 to the middle rings (*R*4 – *R*6) at *Z*_*t*_ = 0. In addition, the shift was more prominent in the case of the *ih-*NPC, and represented a “*handover*” of the tracer from outer rings to inner rings during its translocation through the pore. Further, the handover process appeared to be largely facilitated by the middle FG-Nup rings, *R*4 – *R*6. Although the handover process was observed in both types of pores, the middle rings (*R*4 – *R*6) played a more important role in *ih* −NPC, as illustrated by the prominent peaks in *n*_*b*_ corresponding to the middle rings. The same effects for both types of pores were also observed at *f* = 0.1 for the *hd-*tracer denser patchy tracer (Figure S14). The number of binding contacts near the pore exit (*Z*_*t*_ = −20) were very small because these plots correspond to the trapped tracer trajectories from Figures 2 and 3. Since most of the *hd-*tracer trajectories did not exit the pore, their interactions near the exit (however small) were mostly with the middle rings (*R*4 – *R*6). However, a shift in the *n*_*b*_ plots for *h-*NPC (*blue curves*) toward higher ring indexes can be clearly seen, thus reinforcing the role of the handover process in tracer translocation.

The handover process was also observed for the *sy-*tracer in both *h-*NPC and *ih-*NPC. The *n*_*b*_ plots consistently exhibited shifts toward higher ring indexes as the tracer moved from the entry to the exit in both *h-*NPC and *ih-*NPC (Figure 6B). Since trapping of *sy-*tracer was not observed (see Figures 3D – 3E), *n*_*b*_ values were significant even at the pore exit (*Z*_*t*_ = −20). Similar to the observations in Figure 6A, *n*_*b*_ values at the middle rings were higher for *ih* −NPC compared to *h-* NPC, suggesting a more efficient transfer of the tracer from the pore entry to the exit. Indeed, this was confirmed by the higher *p*_*p*_ value for the *sy-*tracer in *ih-*NPC at *f* = 0.2 (Figure 3F). An interesting interpretation of the *n*_*b*_ plots for distributed patchy tracer in *ih-*NPC is that an efficient *R*4 – *R*6 handover is aided by dynamic binding interactions that can be broken easily as the tracer moves through the pore. For the *sy-*tracer, this is aided by weaker FG-tracer binding interactions compared to those of the *hd-*tracer. This provides a physical justification for setting ɛ_26_ = 1.5 for FG repeats from *R*4 – *R*6 for the trapped trajectories of the denser patchy tracer (Figure 4). As shown in Figure 4, this resulted in the release of all trapped trajectories of the *hd-*tracer, resulting in a very large increase in *p*_*p*_ for *ih-*NPC. Hence, it confirmed the crucial role of the middle rings-facilitated handover process in determining the selectivity of the NPC.

The middle rings-facilitated handover of the denser patchy tracer in the *ih-*NPC is illustrated for a trajectory corresponding to *f* = 0.2 in Figure 7. The time-dependence of *n*_*b*_ for tracer binding with each ring, *R*1 through *R*9, was analyzed during tracer translocation through the *ih-*NPC. Each plot shown in Figure 7 corresponds to tracer binding interactions with a specific FG-Nup ring. The four vertical dashed lines in the plots (*from left to right*) correspond to time instants for; (i) tracer entry into the pore, *Z*_*t*_ = 22.5 (green), (ii) tracer at the mid-plane, *Z*_*t*_ = 0 (black), (iii) decreasing tracer-FG interaction (of middle rings, *R*4 − *R*6) to ɛ_26_ = 1.5 for trapped trajectories (red), and (iv) tracer exit from the pore, *Z*_*t*_ = −22.5 (blue). Though the binding of the peripheral rings (*R*1 – *R*3) to the tracer facilitated its entry into the pore, the middle rings (*R*4 – *R*6) also formed binding contacts with the tracer during the initial entry phase. This is evident from the *n*_*b*_ plots for *R*4 – *R*6 which show significant binding contacts during tracer entry. The large number of binding contacts with *R*4 – *R*6 during entry was a result of reduced *ρ*_*z*_ values for *ih-*NPC (discussed in Figure 5) and observed in the *n*_*b*_ plots of Figure 6. More significantly, *R*4 – *R*6 remained engaged with the tracer till it reached the mid-plane (where it was trapped). After the binding interaction of *R*4 – *R*6 was reduced to ɛ_26_ = 1.5, the tracer was released by *R*4 – *R*6 and handed over to the peripheral rings at the exit, *R*7 – *R*9. Figure S15 of the Supplementary Information illustrates the handover mechanism for a *sy-*tracer in the *ih-*NPC for *f* = 0.2 (for handover mechanism through *h-*NPC, refer Figure S16 and S17). As trapping was not observed for the *sy-*tracer, ɛ_26_ remained constant at 2.0 throughout the simulation. Despite the absence of a simulated tracer release by weakening of ɛ_26_ for *R*4 – *R*6, a handover of the tracer from the middle rings to the peripheral rings *R*7 – *R*9 was observed, resembling the handover process depicted in Figure 7 (Figure S15 and S16).

**Figure 7.**
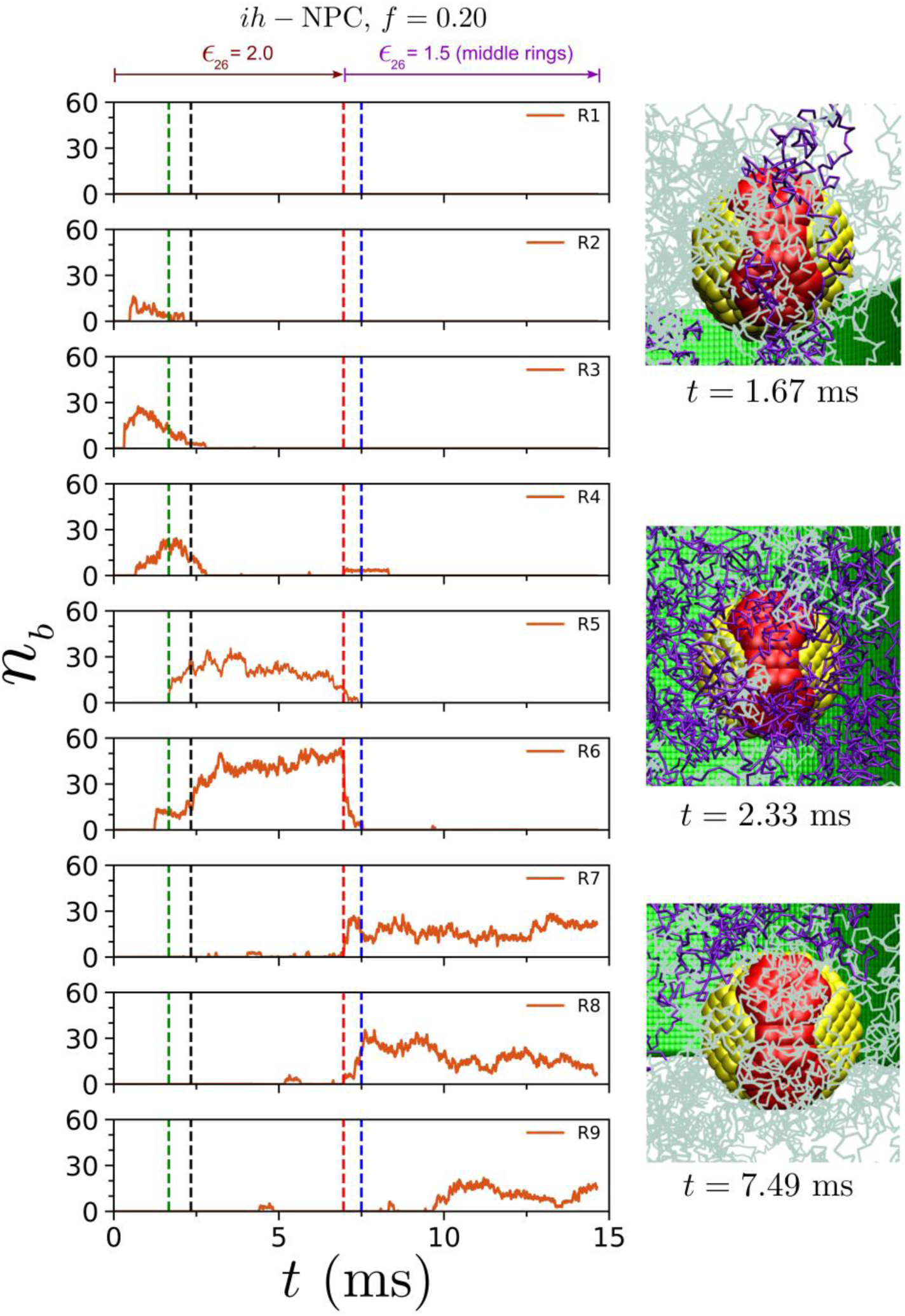
A representative *hd-*tracer trajectory at *f* = 0.2 illustrates the *ring handover* mechanism of enhanced tracer translocation. Plots *n*_*b*_ *versus* time for tracer binding with individual rings show the attachment of different rings to the tracer during the translocation process. During the tracer translocation through the *ih-*NPC, it was sequentially handed over by FG-Nups of successive rings. Simulation snapshots show representative tracer-FG interactions at the entry, mid-plane and exit of the NPC, respectively. Whereas, FG-Nups from the middle three rings are depicted in purple, those from the peripheral rings are shown in light grey.

## CONCLUSIONS

A polymer-physics based minimal model of the NPC and coarse-grained Langevin dynamics simulations were used to elucidate the functional role of FG-Nup polydispersity in NPC transport. The minimal NPC model was a cylindrical pore with a random copolymer brush (comprised of hydrophobic and hydrophilic beads) end-grafted on the inner periphery of the cylinder. The random copolymers represented a sequence-agnostic description of FG-Nups, with hydrophobic segments corresponding to FG repeats in FG-Nups. Three different types of spherical tracers, with and without FG-binding domains, were used to model macromolecular cargo (karyopherin-bound protein cargo) undergoing translocation. Previously, our sequence-agnostic minimal model was successful in capturing both *selectivity* and *specificity* of the NPC, by assuming a monodisperse distribution of FG-Nups along the axial extent of the NPC (23). However, NPCs demonstrate a polydisperse distribution of FG-Nup sizes, with longer FG-Nups occurring near the periphery and shorter FG-Nups confined to the middle portion of the NPC. Hence, tracer transport through a modified, “*inhomogenous”* minimal NPC (*ih-*NPC with longer peripheral FG-Nups and shorter middle FG-Nups) was compared with tracer transport through a *homogeneous* NPC (*h-*NPC with uniform FG-Nup lengths) to investigate the role of FG-Nup polydispersity in NPC transport.

Both types of NPCs rejected the passage of *i*-tracers almost completely; the passage of a small fraction of trajectories being an artefact of our coarse-grained model. In contrast, patchy tracers that modelled karyopherin-bound cargo with either distributed (*sy-*tracer) or localized (*hd-*tracer) binding domains demonstrated selectivity in NPC transport for FG-Nup hydrophobic fractions in the range, 0.1 ≤ *f* ≤ 0.2. Remarkably, this was the same range previously predicted by our minimal NPC model (*h*-NPC) (23), corresponding to the range of FG-repeat fractions found in both *S. Cervisae* and *H. Sapiens*. Significantly, both *sy-*tracer and *hd-*tracer showed increased probability of passage, *p*_*p*_, for the *ih-*NPC (Figures 3 and 4), suggesting a crucial role of the shorter FG-Nup middle rings (*R*4 – *R*6) in enhancing *p*_*p*_ values. For instance, *p*_*p*_ values for an *hd-*tracer through *ih-*NPC were nearly 90% higher compared to those for an *sy-*tracer through *h-*NPC (Figure 4A), suggesting a combined effect of both enhanced karyopherin-FG interactions and FG-Nup length polydispersity. Further, the enhanced selectivity of the *ih-*NPC was demonstrated by high *p*_*e*_ and *p*_*p*_ values (95% and 85%, respectively in Figure 3) for the *hd-*tracer, and the narrower distribution of *p*_*p*_ with respect to *f* (Figure 4A).

Detailed analyses of tracer trajectories through the two NPCs showed that the handover of tracers through successive hydrophobic binding contacts with FG-Nup rings was crucial for successful translocation. Interestingly, tracer handover from rings *R*1 → *R*9 was observed for both *h*-NPC and *ih*-NPC. Without this dynamic binding-unbinding process, tracers were trapped within the pore. Successful translocation was observed primarily within a narrow FG-fraction range (0.1 ≤ *f* ≤ 0.2), consistent with the results of our previous study using the *h-*NPC (23). It was found that the *ring transfer* or *tracer handover* process was more efficient for the *ih*-NPC, with the middle rings *R*4 − *R*6 showing enhanced binding contacts with the tracer. This enhancement in the middle ring binding contacts for the *ih*-NPC (in comparison to *h*-NPC) was true for both *sy-*tracer and *hd-*tracer (Figures 6A and 6B). However, the number of binding contacts were significantly higher for the *hd-*tracer resulting in much higher values of *p*_*p*_.

Hence, both *h*-NPC and *ih*-NPC demonstrated *selectivity* (recognition of tracers) and *specificity* (efficient translocation within a limited FG-fraction range of 0.1 ≤ *f* ≤ 0.2), two features that are characteristic of NPC transport. This range corresponds to the hydrophobic fraction found in humans and yeast (33, 34) NPCs, thus reinforcing the biological relevance of our simulation findings. This study stands out by accurately modeling inhomogeneous pores and denser patchy tracers, closely mimicking the biophysical behavior of NPCs and Kap-cargo complexes. Using a minimal coarse-grained model, we successfully reproduced key aspects of NPC functionality, including the role of shorter FG-Nups in the middle of the pore. Our findings emphasize the importance of brush heterogeneity and provide insights into how the NPC achieves selective and efficient cargo transport. In addition, the results emphasize the evolutionary advantage of varying FG-Nup lengths within NPCs (12, 34, 68, 69). The strategic polydisperse distribution of FG brushes (33, 34) enhances the translocation of Kap-cargo complexes, revealing the biophysical importance of brush heterogeneity in NPC functionality.

## Notes

### Competing Interest Statement

The authors have declared no competing interest.

