## Supplementary Information for "FG-Nup Sequence Length Polydispersity Enhances Selectivity of NPC Translocation"

**Table of Contents** **Page**


| | | |
|---|---|---|
| **Figure S1.** | Inert tracer trajectories from twenty independent simulations through homogeneous pore shown for different values of $f$ | S4 |
| **Figure S2.** | Distributed patchy tracer trajectories from twenty independent simulations through homogeneous pore shown for different values of $f$ ( | S5 |
| **Figure S3.** | Dense patchy tracer trajectories from twenty independent simulations through homogeneous pore shown for different values of $f$ | S6 |
| **Figure S4.** | Inert patchy tracer trajectories from twenty independent simulations through inhomogeneous pore shown for different values of $f$ | S7 |
| **Figure S5.** | Distributed patchy tracer trajectories from twenty independent simulations through inhomogeneous pore shown for different values of $f$ | S8 |
| **Figure S6.** | Dense patchy tracer trajectories from twenty independent simulations through inhomogeneous pore shown for different values of $f$ | S9 |
| **Figure S7.** | Plots showing the equilibrated density distributions of polymer brush segments, considering both hydrophobic and hydrophilic segments. | S10 |
| **Figure S8.** | Plots showing the equilibrated density distributions of polymer brush segments, considering only hydrophilic segments. | S11 |
| **Figure S9.** | Plots showing the equilibrated density distributions of polymer brush segments, considering both hydrophobic and hydrophilic segments. | S12 |
| **Figure S10.** | Plots showing the equilibrated density distributions of polymer brush segments, considering only hydrophilic segments. | S13 |
| **Figure S11.** | Dense patchy tracer trajectories from twenty independent simulations through homogeneous pore shown for different values of $f$ after weakening of tracer-FG hydrophobic affinity. | S14 |
| **Figure S12.** | Dense patchy tracer trajectories from twenty independent simulations through inhomogeneous pore shown for different values of $f$ after weakening of tracer-FG hydrophobic affinity. | S15 |
| **Figure S13.** | Final probabilities associated with different tracer translocation as function of $f$ after weakening hydrophobic interaction between *hd*-tracer and middle rings. | S16 |
| **Figure S14.** | Average number of binding contacts, $\langle n_b \rangle$, between tracer binding domains and FG-repeats corresponding to individual rings, $R1 - R9$, shown for both *h*-NPC and *ih*-NPC at $f = 0.1$ and $f = 0.2$. | S17 |
| **Figure S15.** | Hydrophobic binding contacts between the $sy$ −tracer (distributed patchy tracer) and FG-Nups at $f = 0.2$, during its passage through the *ih*-NPC (inhomogeneous pore). | S18 |

**Figure S16.** Hydrophobic binding contacts between the *sy*-tracer (distributed patchy tracer) and FG-Nups at $f = 0.2$, during its passage through the *h*-NPC (homogeneous pore). S19
**Figure S17.** Hydrophobic binding contacts between the *hd*-tracer (denser patchy tracer) and FG-Nups at $f = 0.2$, during its passage through the *h*-NPC (homogeneous pore). S20

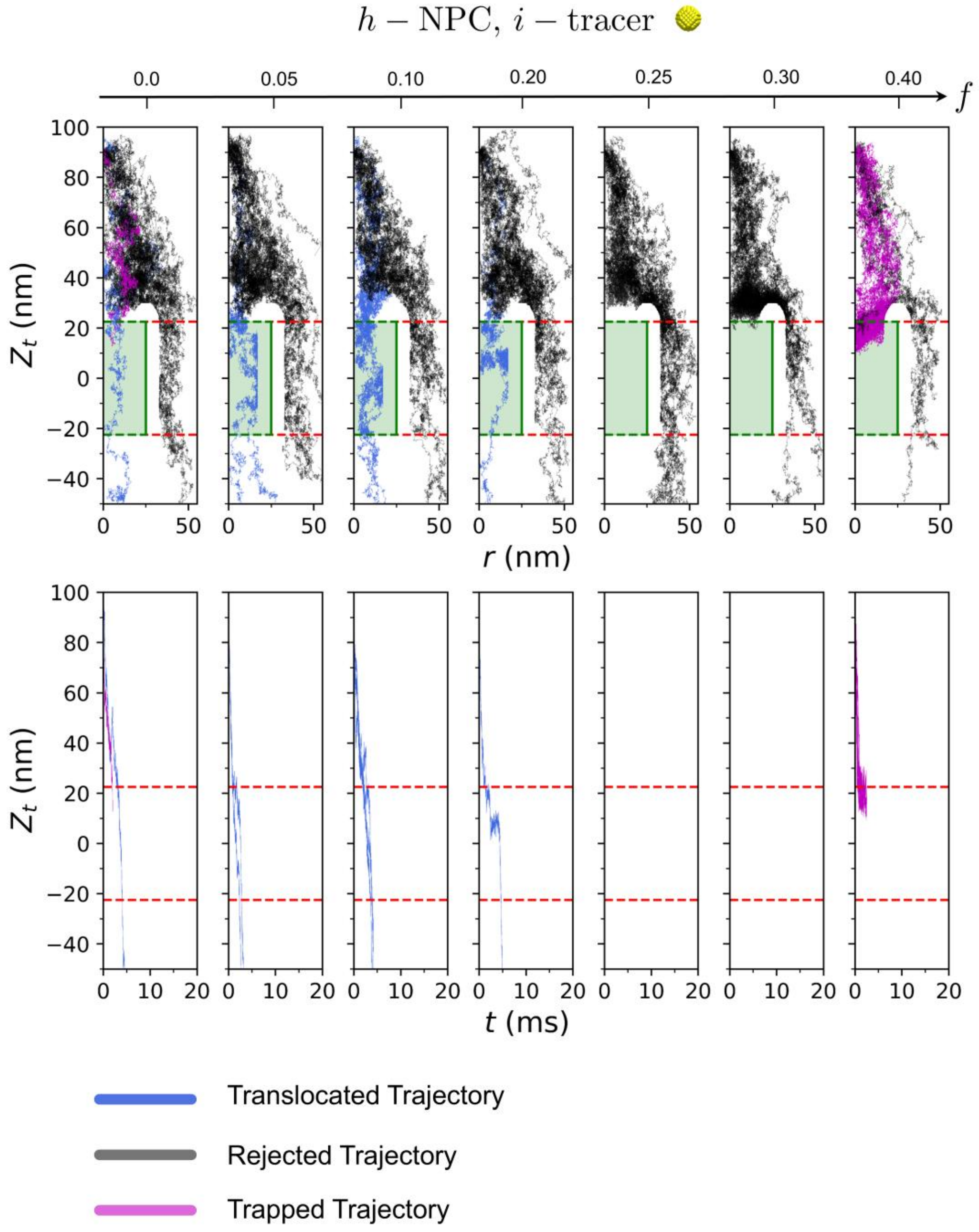


**Figure S1.** Inert tracer trajectories from twenty independent simulations through homogeneous pore shown for different values of $f$ (A) Tracer paths during the simulations represented as $Z_t$ $vs$ $r$ plots, where $Z_t$ is the $z$-coordinate of the tracer, and $r$ is tracer radial coordinate. (B) Plots showing the variation of $Z_t$ with time $t$.

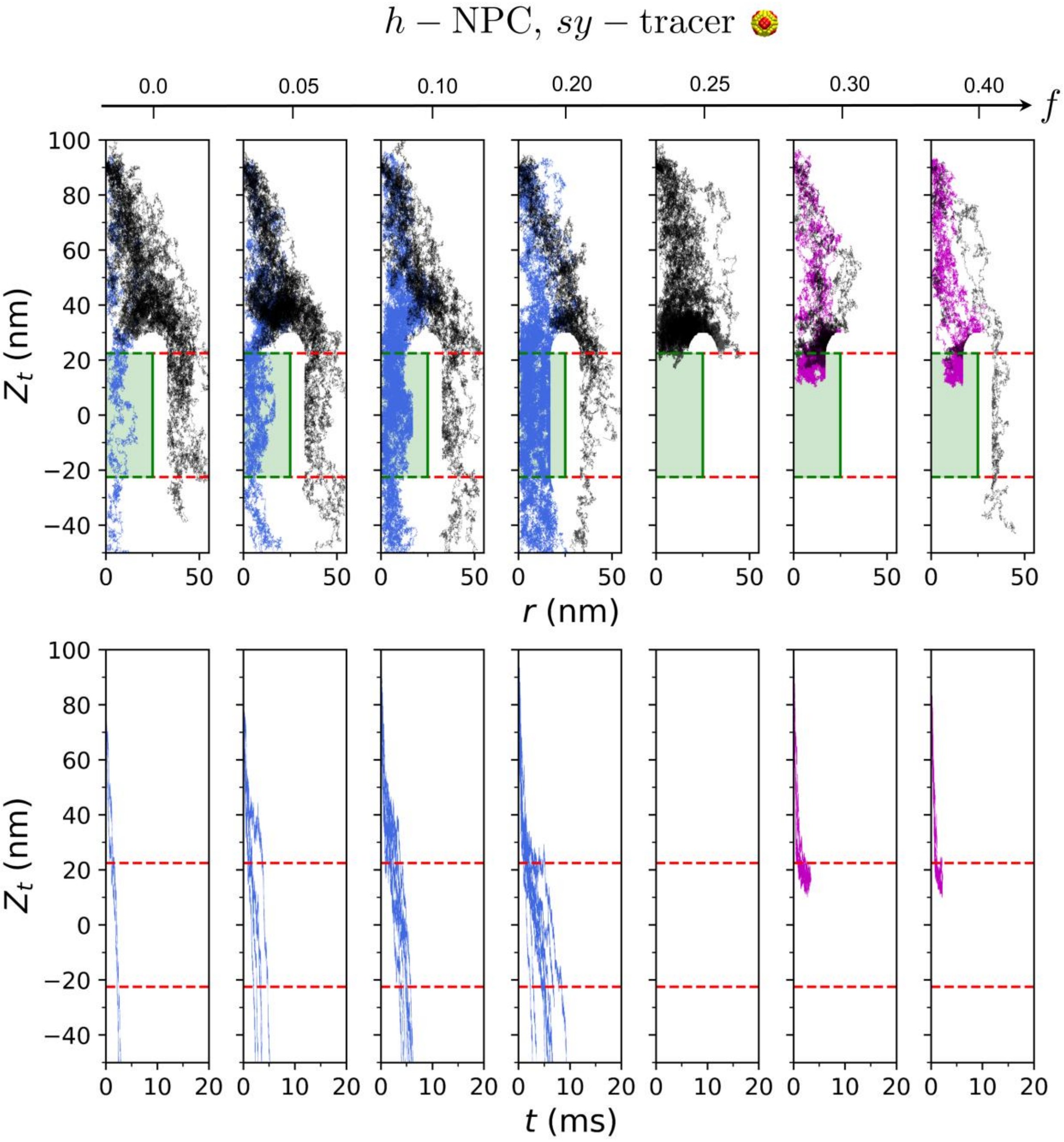


**Figure S2.** Distributed patchy tracer trajectories from twenty independent simulations through homogeneous pore shown for different values of $f$ (A) Tracer paths during the simulations represented as $Z_t$ $vs$ $r$ plots, where $Z_t$ is the $z$-coordinate of the tracer, and $r$ is tracer radial coordinate. (B) Plots showing the variation of $Z_t$ with time $t$.

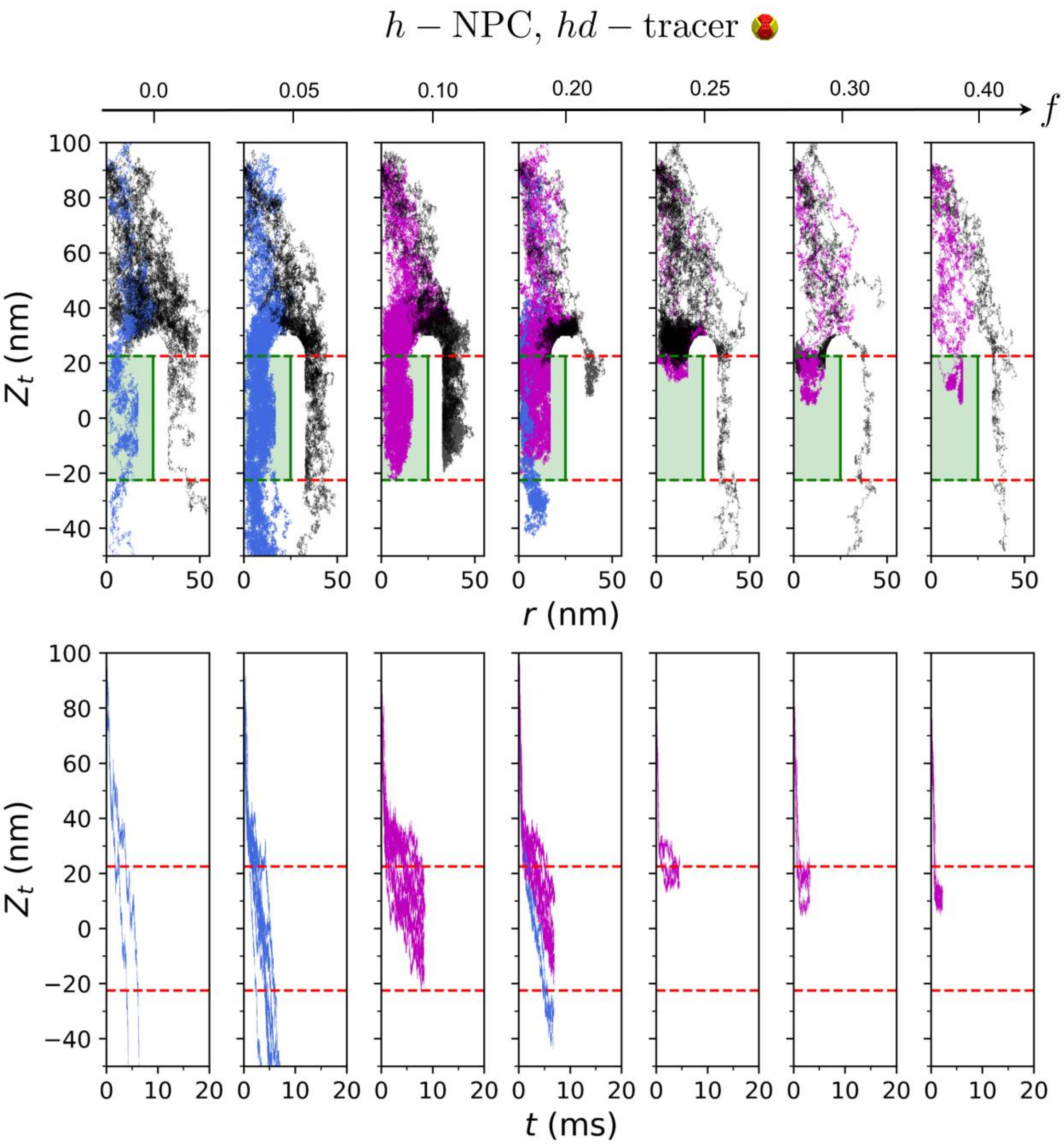


**Figure S3.** Dense patchy tracer trajectories from twenty independent simulations through homogeneous pore shown for different values of $f$ (A) Tracer paths during the simulations represented as $Z_t$ $vs$ $r$ plots, where $Z_t$ is the $z$-coordinate of the tracer, and $r$ is tracer radial coordinate. (B) Plots showing the variation of $Z_t$ with time $t$.

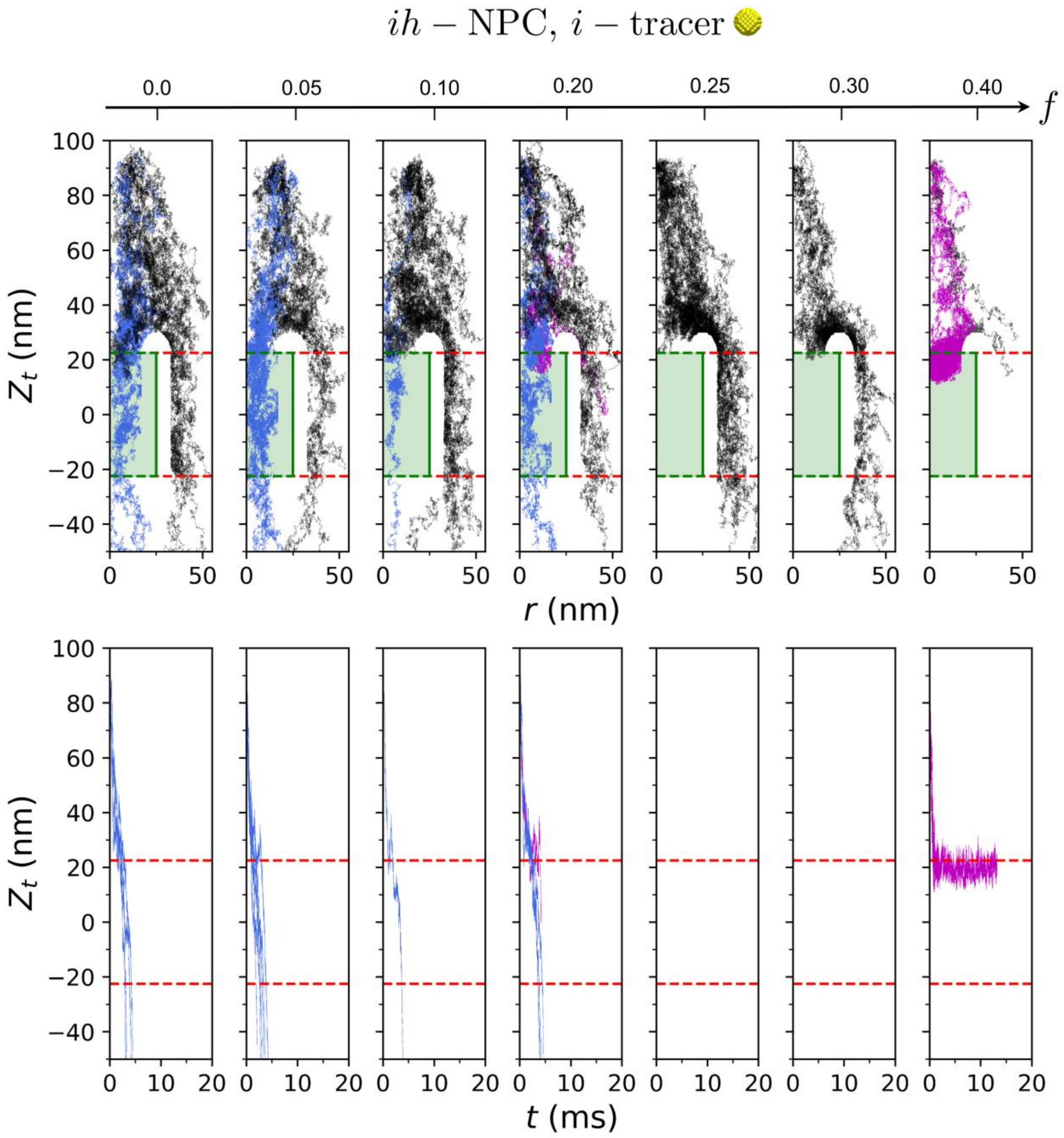


**Figure S4.** Inert patchy tracer trajectories from twenty independent simulations through inhomogeneous pore shown for different values of $f$ (A) Tracer paths during the simulations represented as $Z_t$ $vs$ $r$ plots, where $Z_t$ is the $z$-coordinate of the tracer, and $r$ is tracer radial coordinate. (B) Plots showing the variation of $Z_t$ with time $t$.

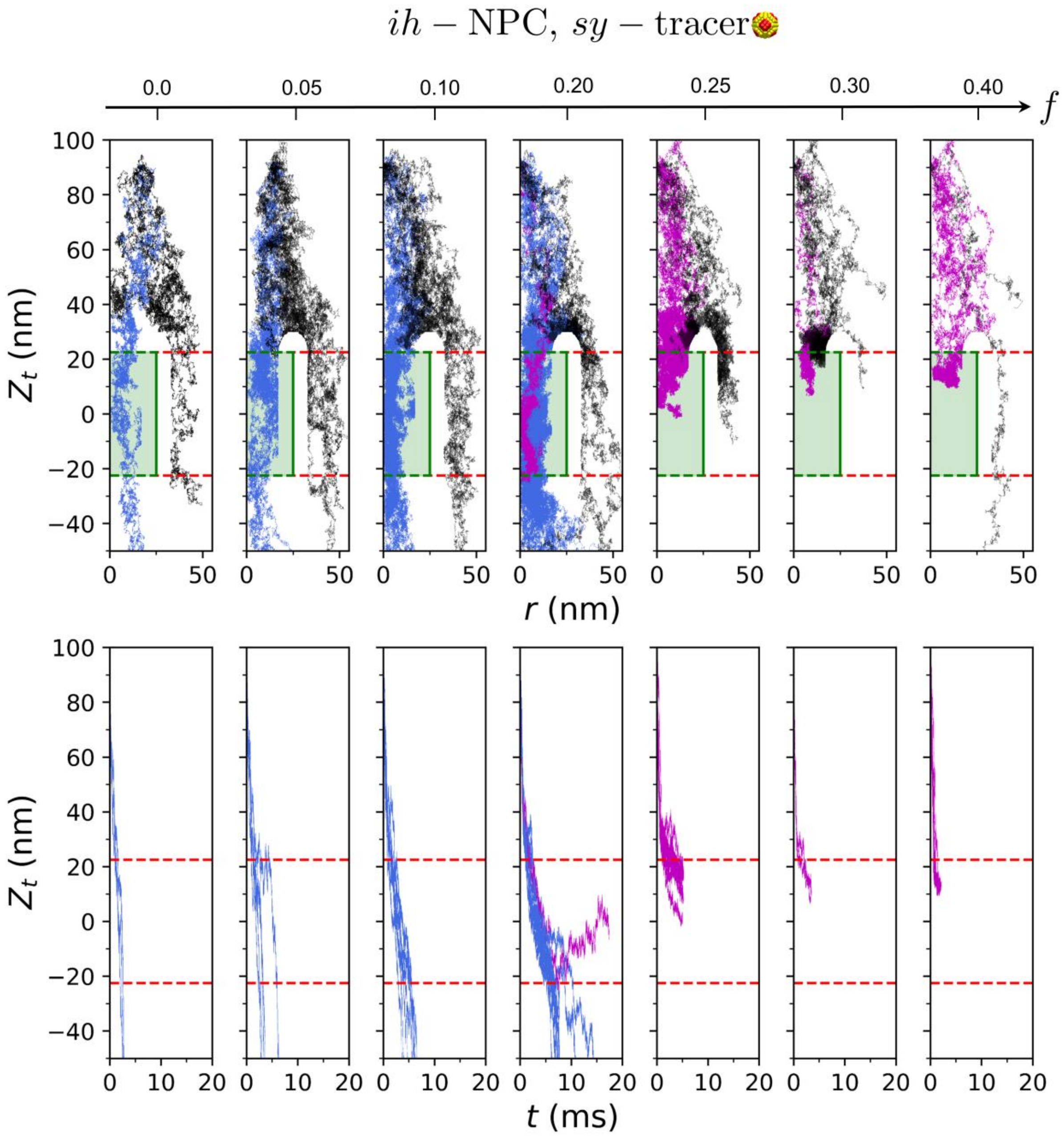


**Figure S5.** Distributed patchy tracer trajectories from twenty independent simulations through inhomogeneous pore shown for different values of $f$ (A) Tracer paths during the simulations represented as $Z_t$ $vs$ $r$ plots, where $Z_t$ is the $z$-coordinate of the tracer, and $r$ is tracer radial coordinate. (B) Plots showing the variation of $Z_t$ with time $t$.

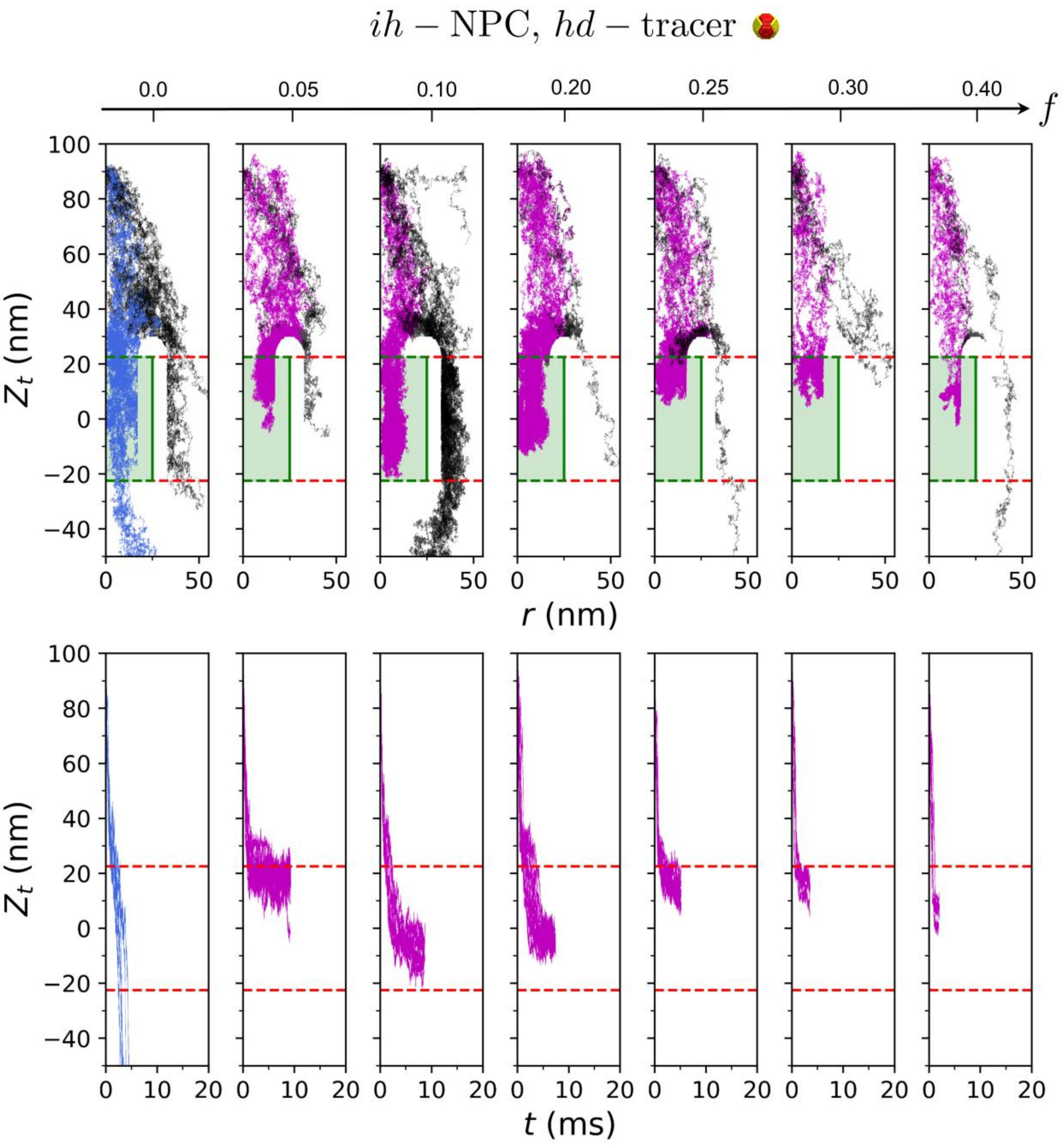


**Figure S6.** Dense patchy tracer trajectories from twenty independent simulations through inhomogeneous pore shown for different values of $f$ (A) Tracer paths during the simulations represented as $Z_t$ $vs$ $r$ plots, where $Z_t$ is the $z$-coordinate of the tracer, and $r$ is tracer radial coordinate. (B) Plots showing the variation of $Z_t$ with time $t$.

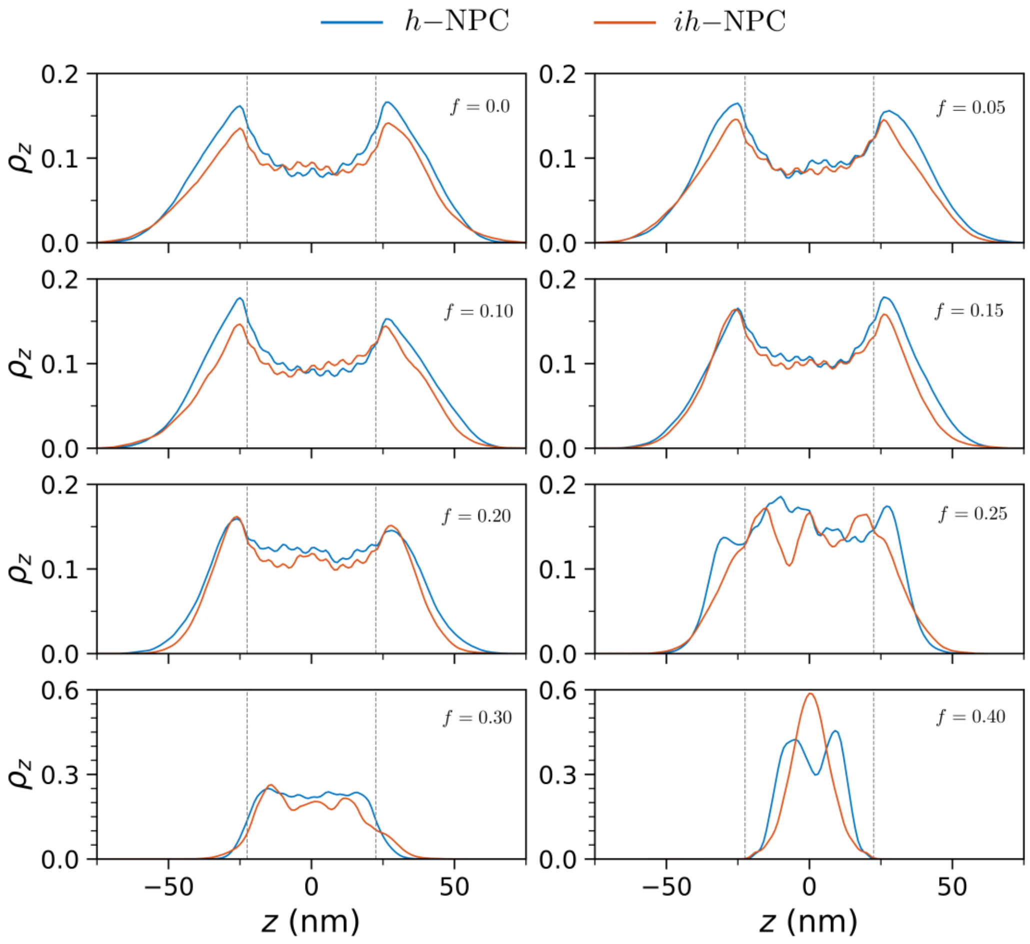


**Figure S7.** Plots showing the equilibrated density distributions of polymer brush segments, considering both hydrophobic and hydrophilic segments. The density distributions are calculated for the full pore by considering the all nine rings of the polymer brushes, plotted along the axial direction of the NPC.

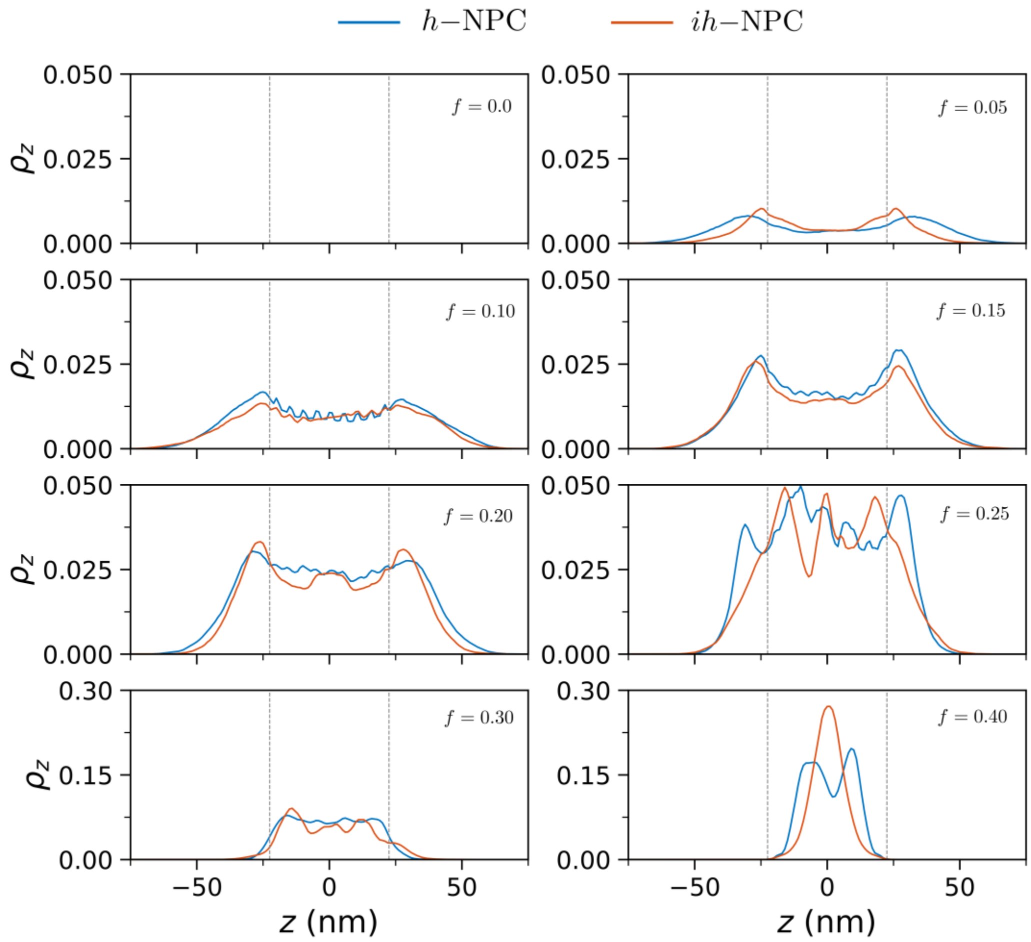


**Figure S8.** Plots showing the equilibrated density distributions of polymer brush segments, considering only hydrophilic segments. The density distributions are calculated for the full pore by considering the all nine rings of the polymer brushes, plotted along the axial direction of the NPC.

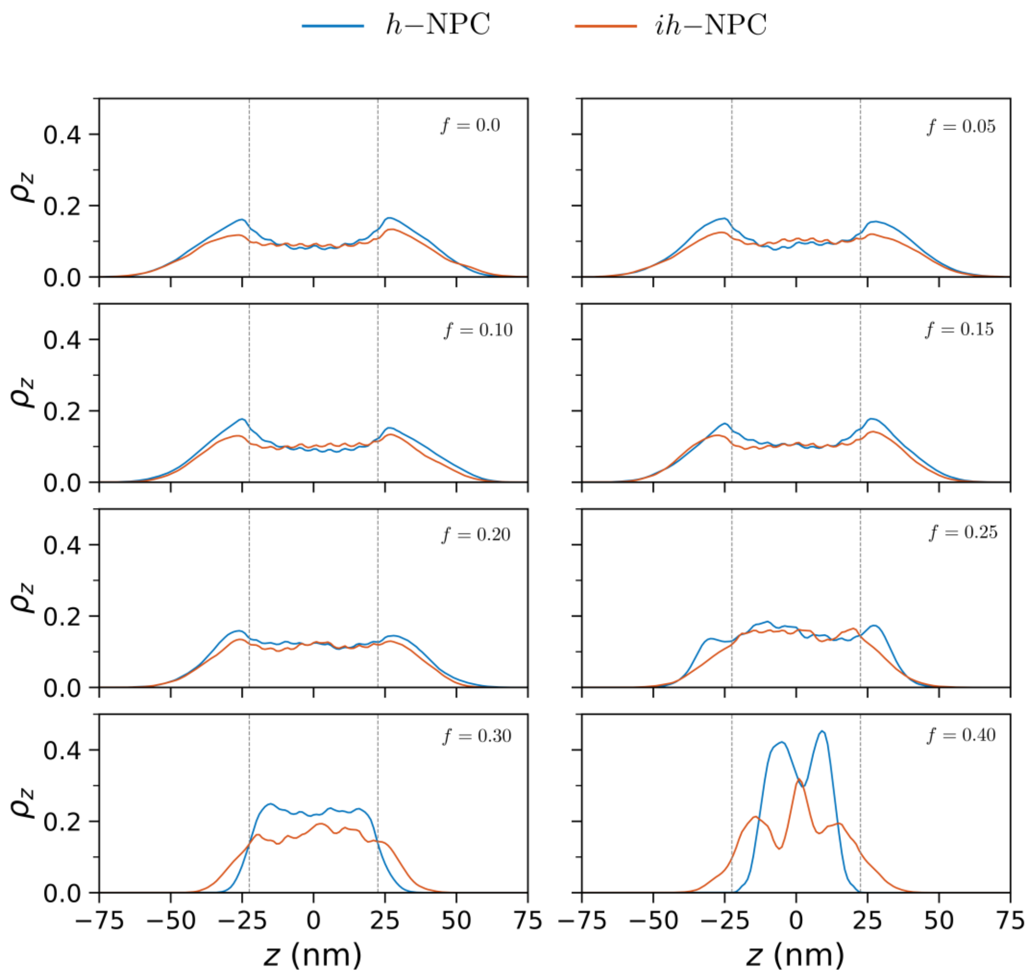


**Figure S9.** Plots showing the equilibrated density distributions of polymer brush segments, considering both hydrophobic and hydrophilic segments. The density distributions are calculated by considering only the middle three rings of the polymer brushes, plotted along the axial direction of the NPC.

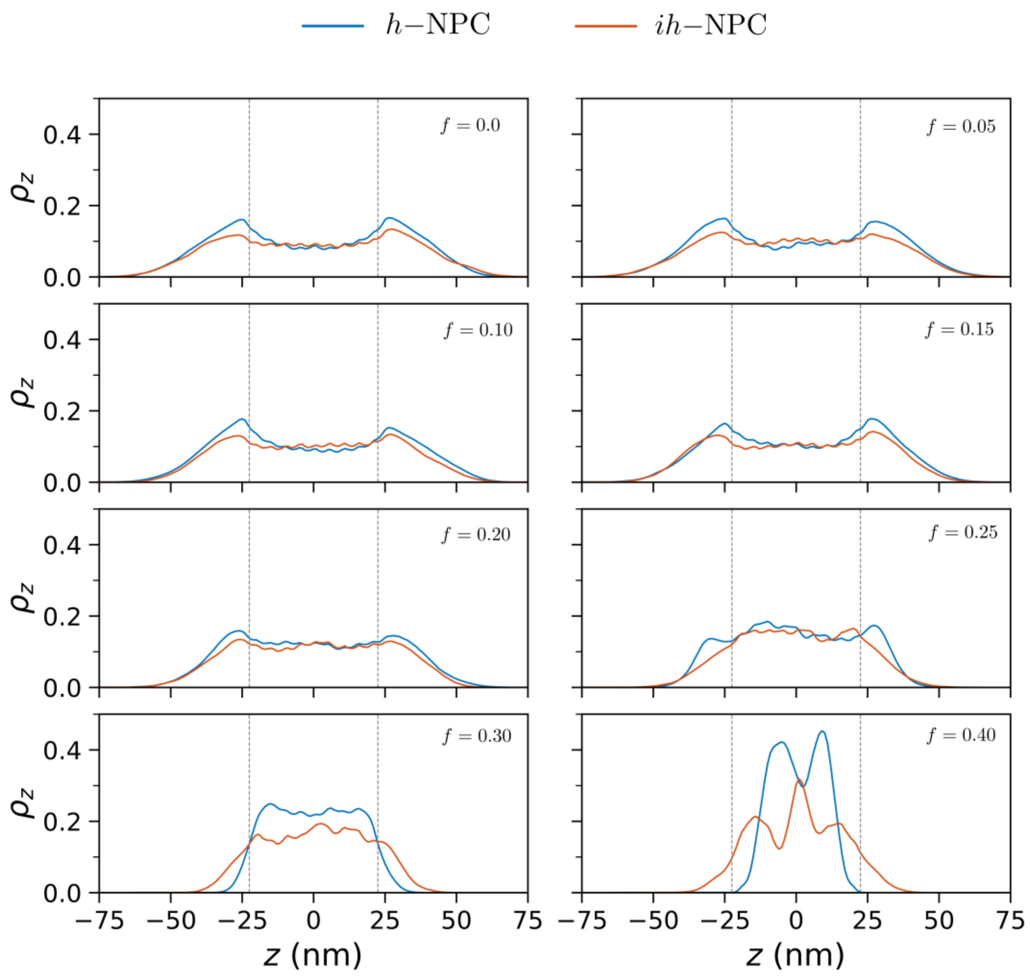


**Figure S10.** Plots showing the equilibrated density distributions of polymer brush segments, considering only hydrophilic segments. The density distributions are calculated for the full pore by considering only the middle three rings of the polymer brushes, plotted along the axial direction of the NPC.

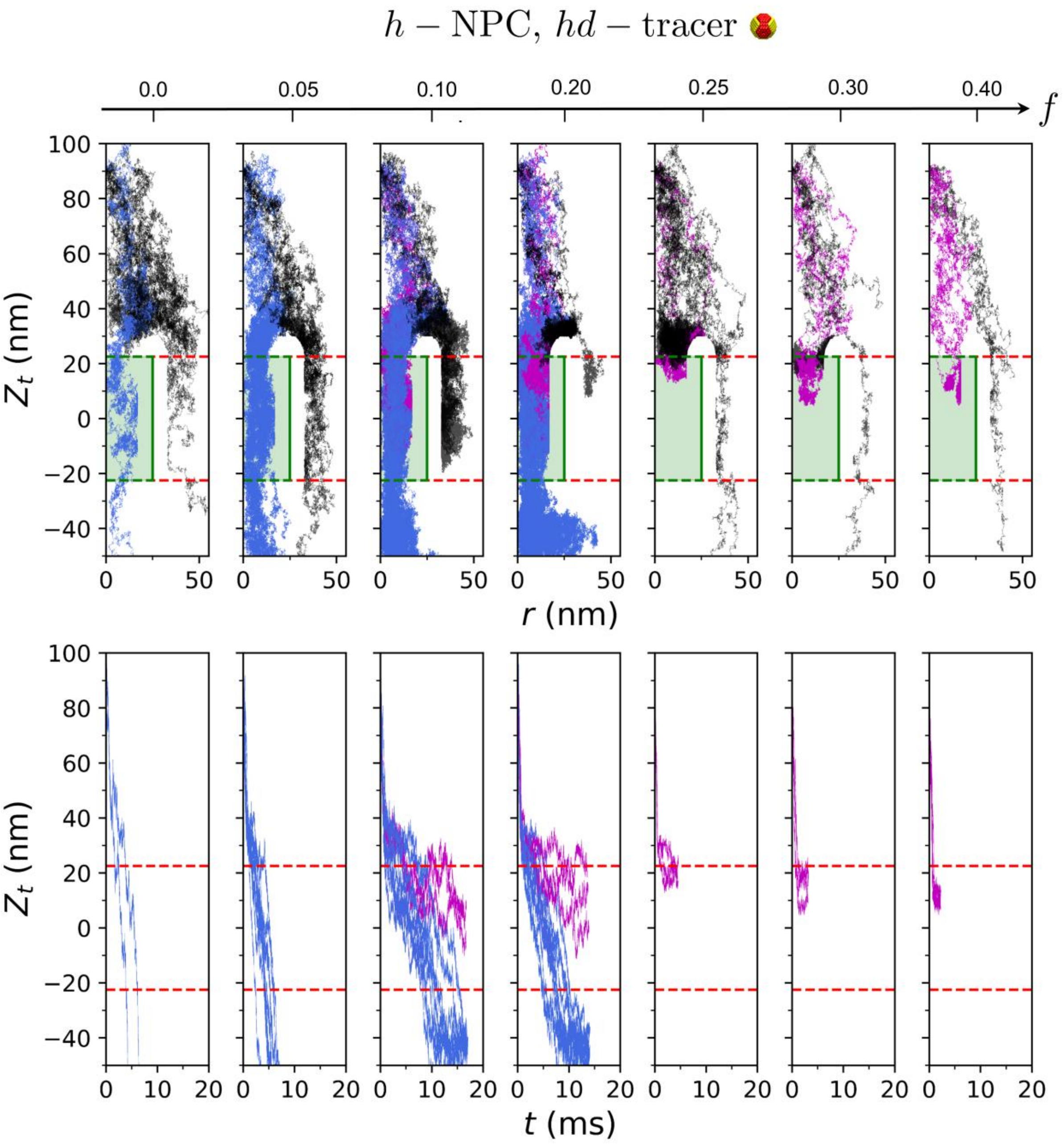


**Figure S11.** Dense patchy tracer trajectories from twenty independent simulations through homogeneous pore shown for different values of $f$ after weakening of tracer-FG hydrophobic affinity (A) Tracer paths during the simulations represented as $Z_t$ $vs$ $r$ plots, where $Z_t$ is the $z$-coordinate of the tracer, and $r$ is tracer radial coordinate. (B) Plots showing the variation of $Z_t$ with time $t$.

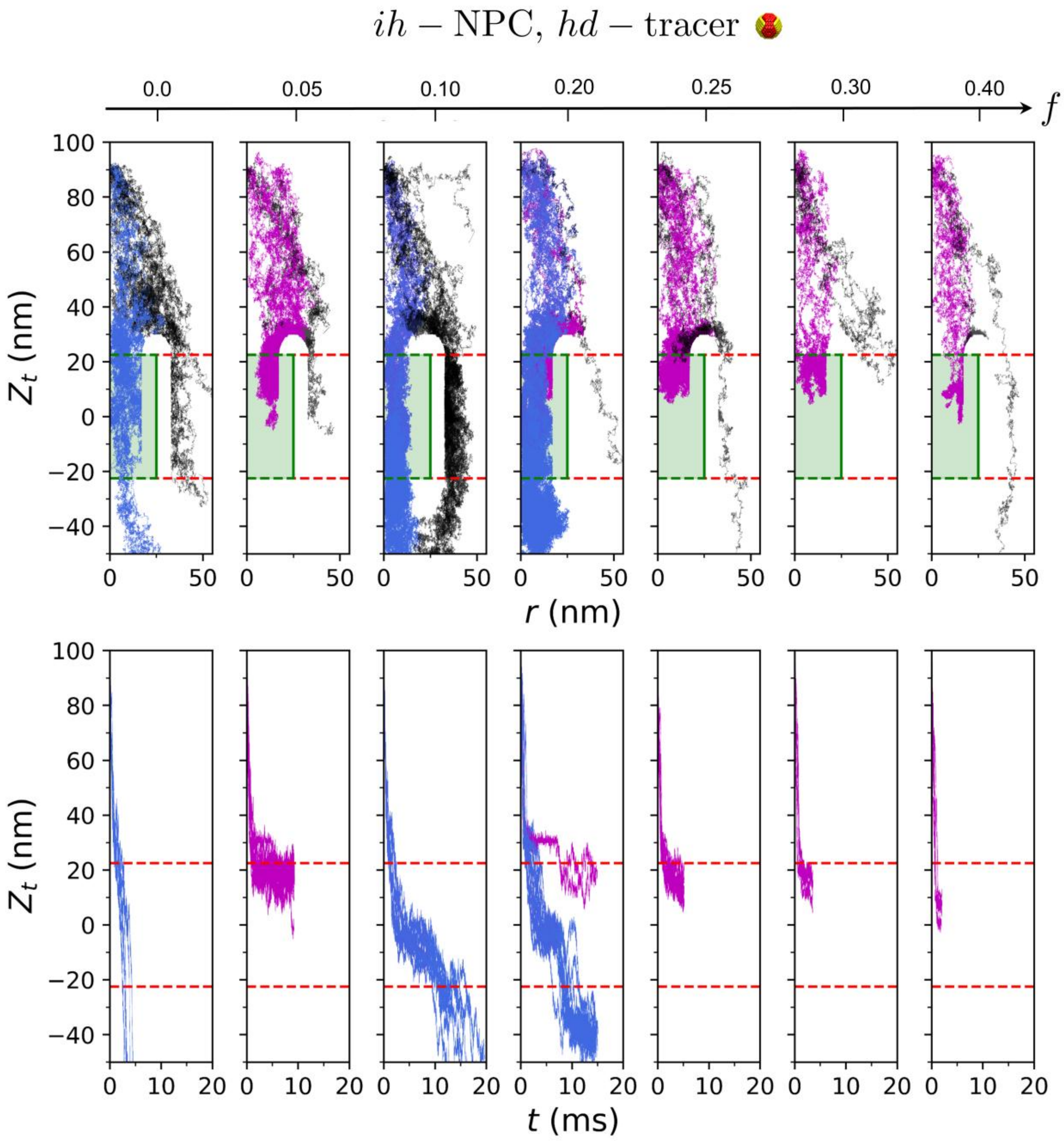


**Figure S12.** Dense patchy tracer trajectories from twenty independent simulations through inhomogeneous pore shown for different values of $f$ after weakening of tracer-FG hydrophobic affinity (A) Tracer paths during the simulations represented as $Z_t$ $vs$ $r$ plots, where $Z_t$ is the $z$-coordinate of the tracer, and $r$ is tracer radial coordinate. (B) Plots showing the variation of $Z_t$ with time $t$.

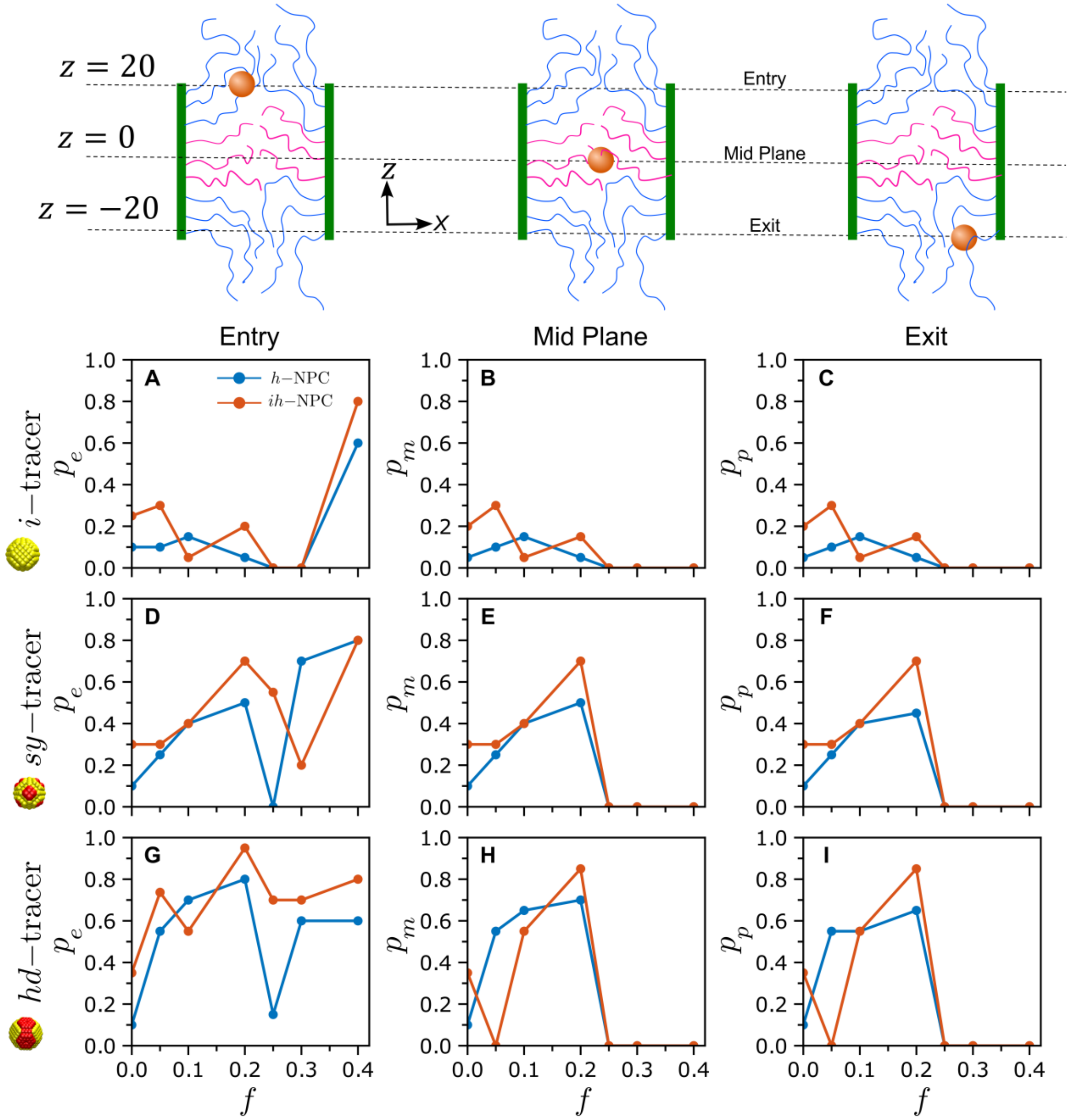


**Figure S13.** Final probabilities associated with different tracer translocation as function of $f$ after weakening hydrophobic interaction between *hd*-tracer and middle rings, including the probability of tracer entry, $p_e$, probability of crossing the NPC mid-plane, $p_m$, and the probability of the tracer completely exiting the pore, $p_p$. Plots are shown for *i*-tracer (A – C), *sy*-tracer (D – F) and *hd*-tracer (G – I) corresponding to both $h$ −NPC and $ih$ −NPC.

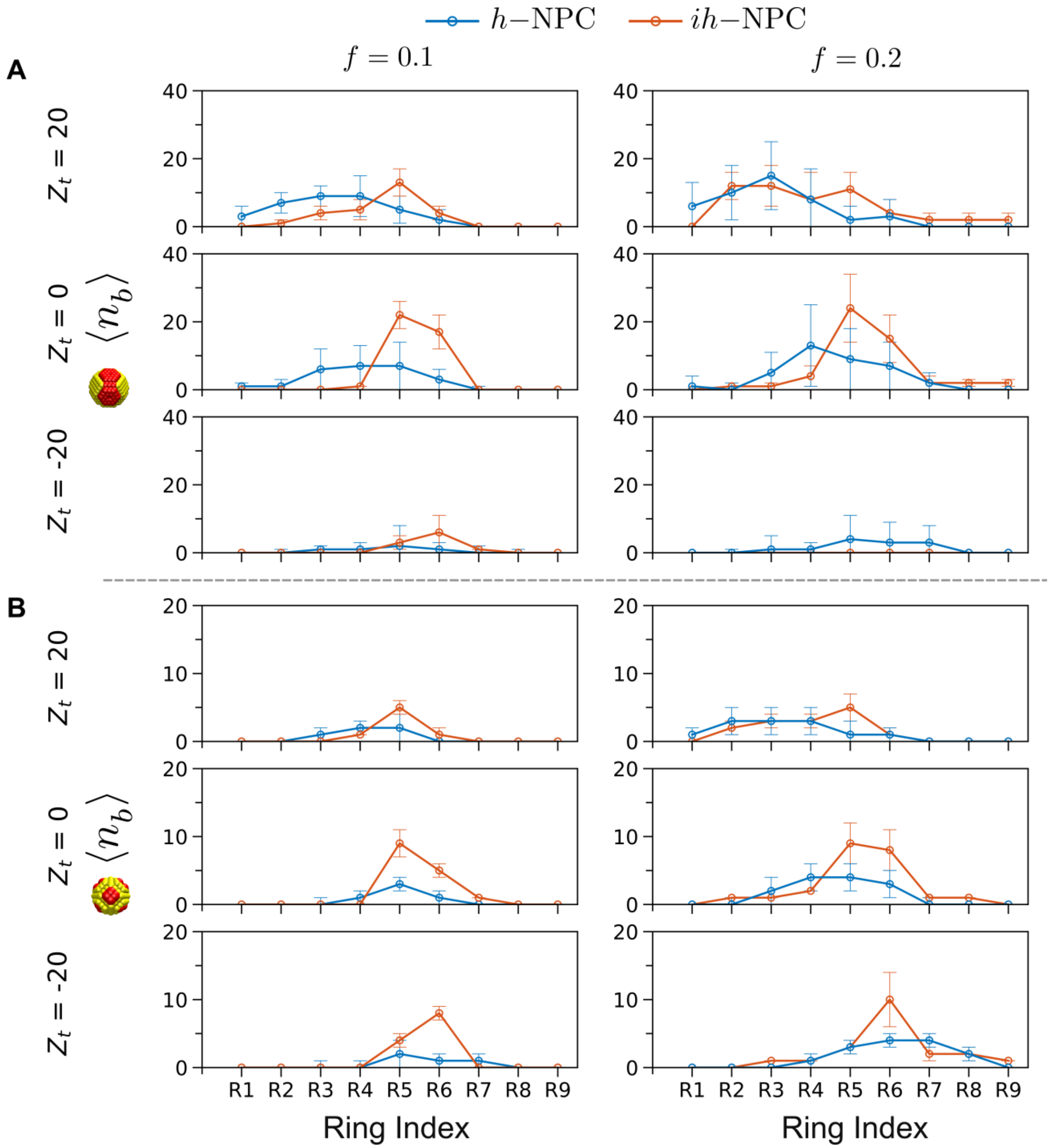


**Figure S14.** Average number of binding contacts, $\langle n_b \rangle$, between tracer binding domains and FG-repeats corresponding to individual rings, $R1 - R9$, shown for both *h*-NPC and *ih*-NPC at $f = 0.1$ and $f = 0.2$. The variation in $\langle n_b \rangle$ with respect to the ring index are shown at the entry ($Z_t = 20$), mid-plane ($Z_t = 0$) and exit ($Z_t = -20$) of the NPCs for (A) *hd*-tracer and (B) *sy*-tracer.

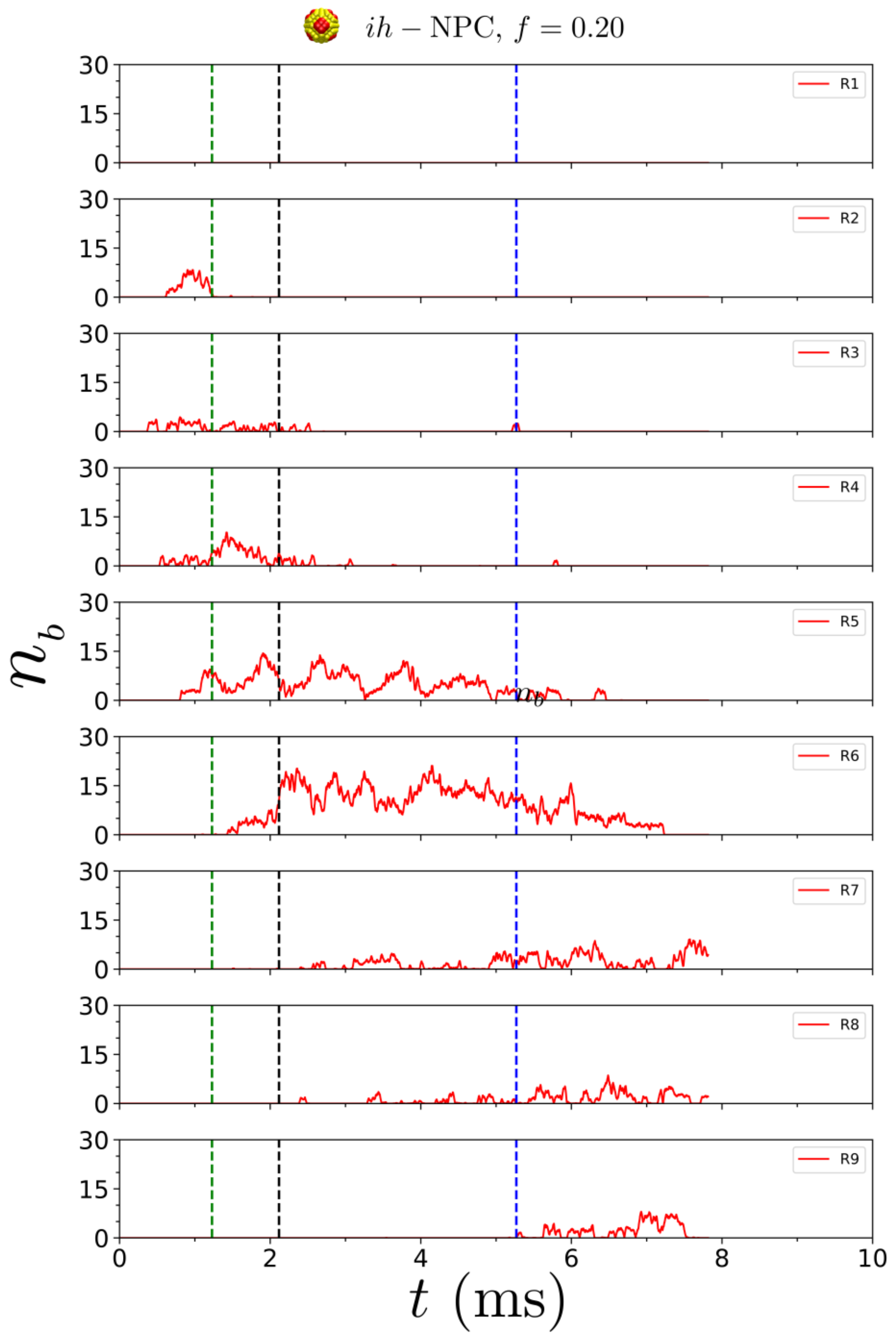


**Figure S15.** Hydrophobic binding contacts between the *sy*-tracer (distributed patchy tracer) and FG-Nups at $f = 0.2$, during its passage through the *ih*-NPC (inhomogeneous pore). The tracer is sequentially handed over from one FG-Nup to another as it enters and moves downwards through the pore.

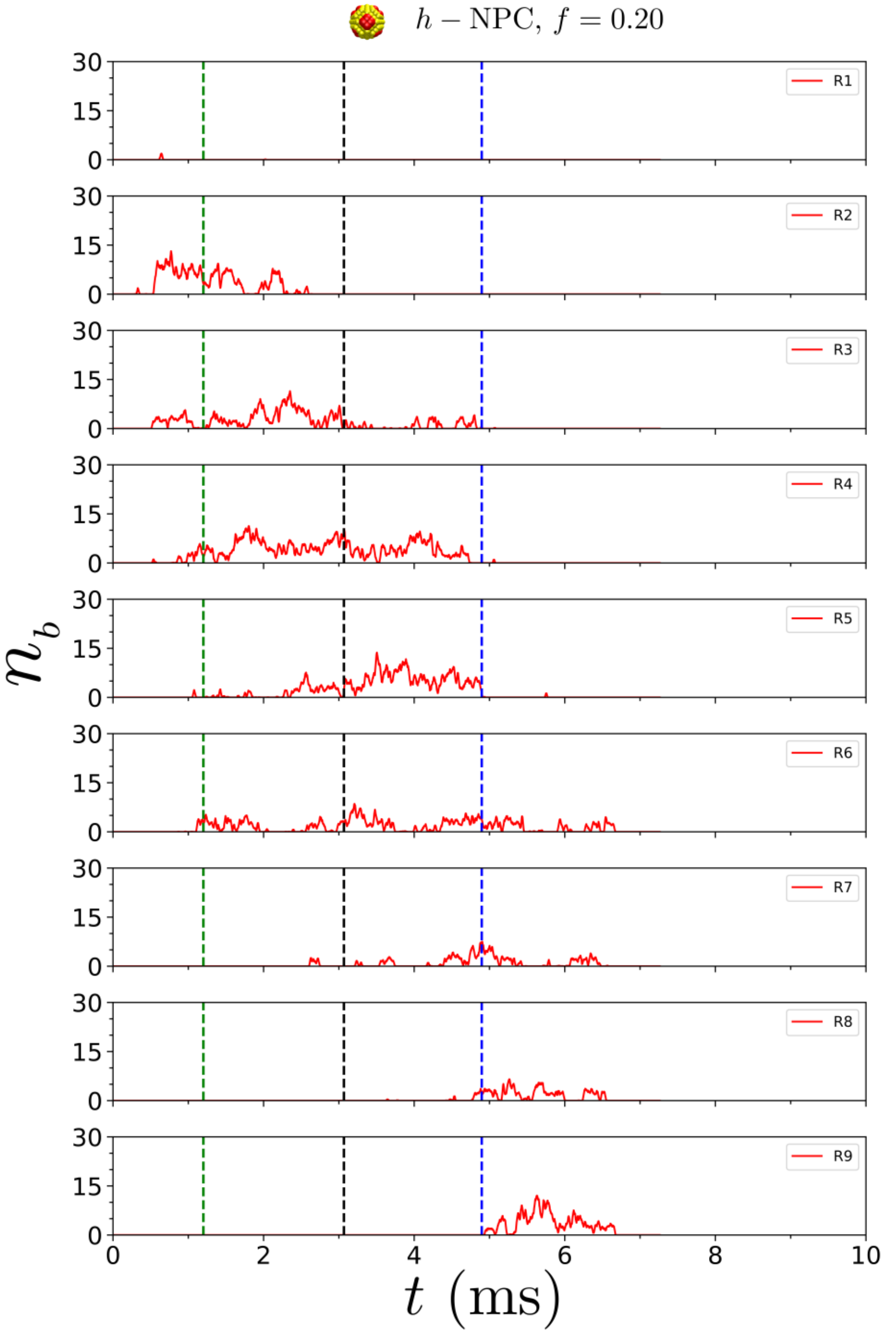


**Figure S16.** Hydrophobic binding contacts between the *sy*-tracer (distributed patchy tracer) and FG-Nups at $f = 0.2$, during its passage through the *h*-NPC (homogeneous pore). The tracer is sequentially handed over from one FG-Nup to another as it enters and moves downwards through the pore.

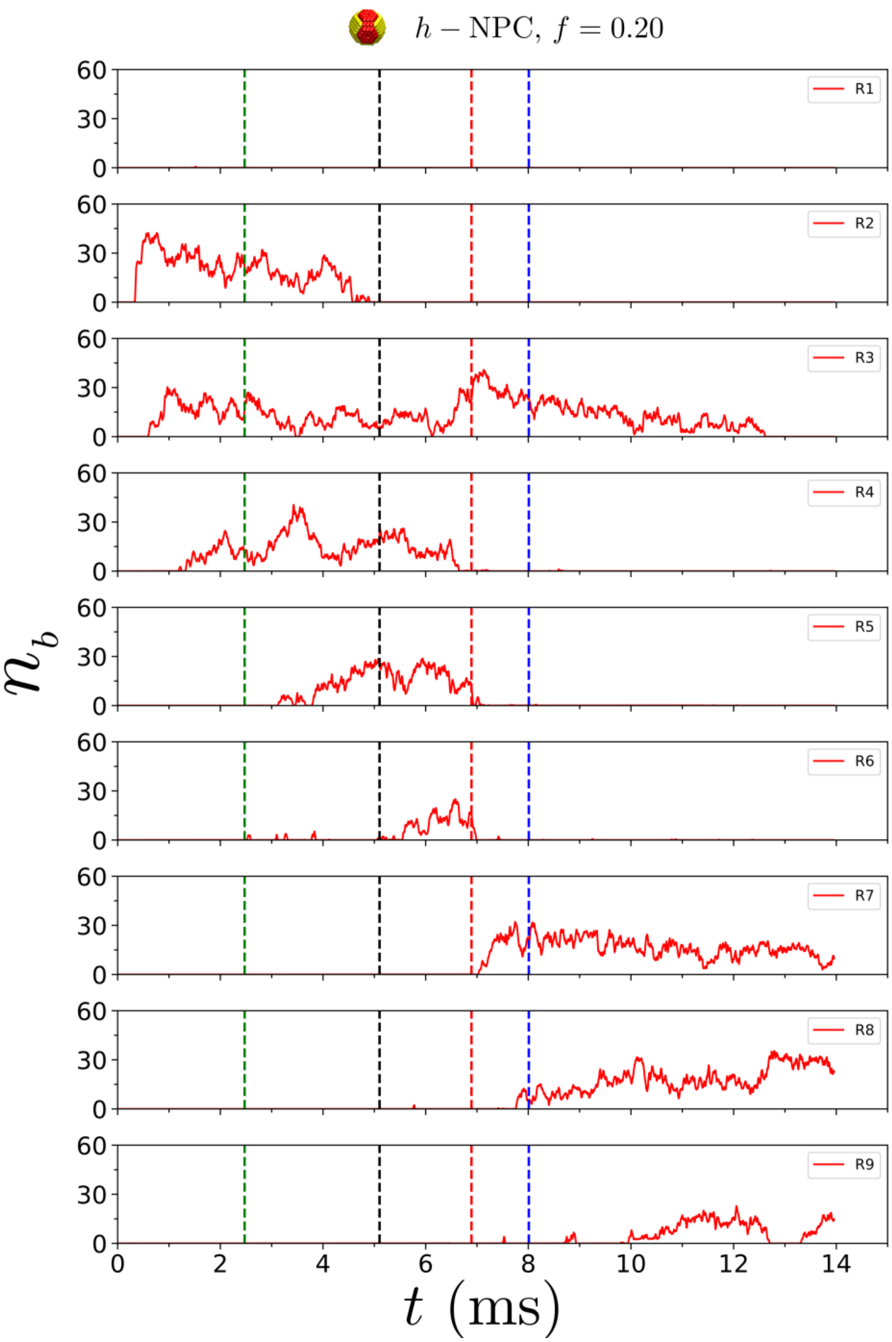


**Figure S17.** Hydrophobic binding contacts between the *hd*-tracer (denser patchy tracer) and FG-Nups at $f = 0.2$, during its passage through the *h*-NPC (homogeneous pore). The tracer is sequentially handed over from one FG-Nup to another as it enters and moves downwards through the pore.
